# A Translational Reference for Green Autofluorescence Imaging in the Rhesus Macaque Eye using the OcuMet Beacon

**DOI:** 10.64898/2026.08.11.744247

**Authors:** Ana Ripolles-Garcia, Jaegook Lim, Ana C Raposo, J’adore C Bailey, Karin W Handel, Meher J Khan, Lindsey R Sutton, Jennifer Yu, Emily K Dougherty, Tracy Nguyen Jaggers, Brian Lam, Yasmine N Valjalo, Rosie Thienpaitoon, Nathaly A Muniz, Elizabeth Giorgi, Carol Iveth Villafuerte-Trisolini, Karen Anderson, Nadeen Habbas-Nimer, Collin A Rich, Kurt Riegger, Ala Moshiri, Brian C Leonard, Glenn Yiu, Sara M Thomasy

**Affiliations:** Dept. of Surgical & Radiological Sciences, School of Veterinary Medicine, University of California, Davis, Davis, CA; Dept. of Ophthalmology & Vision Science, School of Medicine, University of California, Davis, Sacramento, CA; Stoke Therapeutics Inc, Bedford, MA; OcuSciences Inc., Ann Arbor, MI; National Biomedical Research Institute, University of California, Davis, Davis, CA

**Keywords:** Green autofluorescence, Retinal imaging, Oxidative stress, Retinal metabolic imaging, Rhesus macaque

## Abstract

**Purpose:** To evaluate associations between green autofluorescence (GAF) and structural and functional measures relevant to retinal and optic neuropathies, and to establish normative GAF values across the optic nerve head (ONH), macula, and papillofoveal bundle (PFB) in rhesus macaques.

**Methods:** Eighty-two macaques with normal ONH morphology by spectral-domain optical coherence tomography (SD-OCT) were included with a mean ± SD age of 12.76 ± 7.18 (range 0.11-29.39) years. The GAF images were acquired in the ONH, macula and PFB with the OcuMet Beacon. In a subset of macaques (n=19), pattern electroretinogram (PERG) and photopic full-field ERG including the photopic negative response (PhNR) were recorded.

**Results:** The GAF significantly increased with age in the ONH, macula and PFB. After adjusting by age, there were no sex differences, but IOP showed a positive association with macular GAF. At the ONH, higher GAF correlated with thinner retinal nerve fiber layer, inner and outer segment complex, and total retinal thickness. In the macula, inner nuclear layer thickness was positively associated with GAF, whereas outer plexiform layer and inner and outer segment complex were inversely associated. The PERG amplitudes inversely tracked ONH GAF.

**Conclusions:** GAF rises with age and IOP, couples to retinal structure, and at the ONH, aligns with inner-retinal functional indices. This study provides a regional reference for GAF in rhesus macaques.

**Translational Relevance:** Normative GAF data in healthy rhesus macaques provide a framework for interpreting this noninvasive signal in translational studies of retinal and optic nerve disease.

## INTRODUCTION

Among ocular tissues, the retina and optic nerve head (ONH) are uniquely energy demanding and supported by extensive mitochondrial activity.^1^ Subtle disturbances in redox balance here can precede structural change, underscoring the need for a sensitive, noninvasive assessment of mitochondrial function.^2^ Mitochondria, being primary sites of reactive oxygen species (ROS) production, are particularly susceptible to oxidative stress, which can impair their function and contribute to cell dysfunction.^3^ Flavoproteins are a class of proteins that contain a flavin cofactor, such as flavin adenine dinucleotide or flavin mononucleotide, which play essential roles in cellular redox reactions, including mitochondrial electron transport and energy production.^4^ These proteins are involved in oxidative metabolism, and their fluorescence properties allow for direct visualization of mitochondrial function. Flavoprotein fluorescence is an intrinsic green light (520–540 nm) emitted by oxidized mitochondrial flavoproteins when excited by a blue spectrum light (455–470 nm), because the isoalloxazine ring absorbs blue light and then releases green fluorescence, whereas in the reduced state the excited energy is dissipated without light emission, making it a sensitive marker of oxidative stress and mitochondrial dysfunction.^5^ Therefore, noninvasive *in vivo* imaging of green autofluorescence (GAF) using specialized cameras may provide indirect insight into retinal metabolic activity and oxidative stress.^6^ However, because *in vivo* ocular imaging cannot definitively isolate flavoprotein fluorescence from other endogenous fluorophores with overlapping excitation and emission spectra, the measured signal should not be interpreted as a fluorophore-specific measurement of mitochondrial flavoprotein fluorescence. Therefore, throughout this manuscript, we refer to the measured output as GAF.

Elevated GAF concentrations are observed in mitochondrial retinopathies, a group of disorders characterized by mitochondrial dysfunction, including diabetic retinopathy and age-related macular degeneration, where oxidative damage contributes to disease progression.^5,7,8^ Similarly, optic neuropathies such as glaucoma and Leber’s hereditary optic neuropathy exhibit elevated GAF, reflecting compromised mitochondrial function in retinal ganglion cells and the ONH.^6,9,10^ Importantly, GAF serves as an early indicator of cellular dysfunction, which is critical to enable rapid intervention and widen the therapeutic window to preserve vision.^6,9,11^ This capability enhances the utility of GAF as a diagnostic tool and sensitive monitoring of disease progression or response to treatment but it should be first evaluated in healthy populations to determine its variability within and between individuals.^12^

Rhesus macaques (*Macaca mulatta*) are widely used as a large animal model for studying retinal and optic nerve diseases due to their close anatomical and physiological resemblance to humans.^13^ Their foveated retina, similar eye size, and susceptibility to comparable pathophysiological changes make them an invaluable model for ophthalmic research.^13^ Establishing normative GAF data in these rhesus macaques is essential for distinguishing between normal metabolic variations and disease-induced changes. In addition, GAF can complement the characterization of established nonhuman primate disease paradigms, including inducible glaucoma models,^14–16^ as well as genetically defined, spontaneous optic neuropathy models such as *OPA1* associated autosomal dominant optic atrophy identified in rhesus macaques.^17^ Given that age-related changes in the ONH and retina have been documented in both species, developing a well-defined baseline for GAF in rhesus macaques will aid in interpreting variations associated with aging or disease progression.^18,19^ Furthermore, GAF in rhesus macaques may facilitate the early detection of responses to experimental therapies in diseases characterized by slow progression. In such cases, assessing disease stabilization using conventional structural and functional methods, such as optical coherence tomography (OCT) or electroretinography (ERG), would require extended and cost-intensive follow-up periods due to the delayed manifestation of structural and functional changes. To our knowledge, no comprehensive or large-scale GAF fundus imaging studies in humans, rhesus macaques or other animal models have been published, while clinical reports are limited to small human populations and pilot studies.^9,20,21^

The primary objective of this study was to determine if GAF intensity across three anatomical regions, ONH, macula, and papillofoveal bundle (PFB), correlates with established structural and functional parameters relevant to retinal and optic nerve health. A secondary goal was to establish a normative database of GAF in healthy rhesus macaques for the ONH, macula, and PFB. By developing this reference dataset, we aimed to improve understanding of the distribution and behavior of the recorded GAF signal in healthy rhesus macaques and to provide a framework for its interpretation in future studies. We hypothesized that GAF measured in healthy rhesus macaques would vary systematically with age and retinal region, and that these measurements would show quantifiable associations with ocular structural and functional parameters.

## MATERIAL AND METHODS

### Rhesus macaque subjects and housing

All animals used in this study were rhesus macaques (*Macaca mulatta*) housed at the National Biomedical Research Institute (NBRI), which is accredited by the Association for Assessment and Accreditation of Laboratory Animal Care International (AAALAC). Throughout the study, guidelines from the Association for Research in Vision and Ophthalmology (ARVO) Statement for the use of rhesus macaques in Ophthalmic and Vision Research were strictly followed. The study complied with NIH guidelines for the care and use of laboratory rhesus macaques and was approved by the UC Davis Institutional Animal Care and Use Committee.

### Ophthalmic phenotyping

Phenotypic data were collected from normal rhesus macaques during routine prescreening and study intake to confirm the absence of age related or inherited ocular abnormalities. Examinations were performed under sedation or, when required, general anesthesia. Sedation consisted of intramuscular injections of ketamine (5 to 30 mg/kg), midazolam (0.1 mg/kg), and dexmedetomidine (0.05 to 0.075 mg/kg). Mydriasis was achieved with tropicamide (1%; Bausch & Lomb, Bridgewater, NJ) and phenylephrine (2.5%; Paragon Biosciences, Northbrook, IL), and cycloplegia was achieved with cyclopentolate (1%; Akorn, Lake Forest, IL). Complete ocular exams were performed by a boarded veterinary or physician ophthalmologist and included slit lamp biomicroscopy, indirect ophthalmoscopy, fluorescein staining, measurement of intraocular pressure (IOP), fundus photography, and spectral domain optical coherence tomography (SD-OCT) with confocal scanning laser ophthalmoscopy (cSLO; Spectralis; Heidelberg Engineering, Heidelberg, Germany); IOP was measured with rebound tonometry (Icare TonoVet, Icare Finland Oy, Helsinki, Finland).

### Green autofluorescence imaging

Functional imaging of the retina and ONH was performed by quantifying GAF using a recently FDA-cleared device (OcuMet Beacon®, OcuSciences, Ann Arbor, MI). Regions of interest included the ONH, macula, and the PFB. Each acquisition began with a 21.5°(V)x60°(H) infrared reference image, followed by 21.5°(V)x17°(H) GAF imaging. All acquisitions received lens fluorescence compensation. Algorithmic evaluation of image quality automatically selected the highest-contrast image from multiple captures per session, ensuring adequate illumination with minimal pupil cropping.

The OcuMet Beacon incorporates an infrared scanner and a fluorescence detection channel configured to quantify fluorescence in a spectral range expected to be dominated by flavoprotein fluorescence. For each eye and each region of interest at least three short acquisitions were obtained. Frames with motion, blink, registration/segmentation failure, pupil cropping, glare, off-center targets, or other fluorescent contaminants were discarded. Focus, centering, and registration to the infrared reference were verified. A MATLAB image quality index was computed for both infrared and GAF images. This index is scored on a scale from 0 to 100, with scores below 30 considered poor, scores from 30 to <70 considered acceptable, and scores from 70 to 100 considered good. The image-quality index is distinct from the signal-to-noise ratio and is not expressed in decibels. Acquisitions with an index below 30 were excluded. Proprietary software was then used to calculate GAF parameters for each image.

In the ONH region, software algorithms identified the ONH margins on the infrared image and demarcated a 32-sided polygon (32 vertices) around the ONH rim, with inner and outer boundaries corresponding to 0.5-1.0 times the ONH rim size. GAF was quantified within this annulus, which exhibits the highest concentration of GAF signal, consistent with the distribution of mitochondria at the ONH. The annulus was then divided into temporal, superior, nasal, and inferior sectors, with global GAF defined as the average of all four sectors. A GAF rim profile was generated to display GAF levels circumferentially across these anatomic quadrants, aiding in the localization of specific regional changes. In the PFB region, GAF intensity was measured within an ellipsoidal, teardrop-shaped area defined by longitudinal edges at the centers of the fovea and ONH, as it had been previously established.^22^ This area was then further subdivided into the arcuate and horizontal bundles. Additionally, at the ONH, the PFB teardrop was cropped at 1.3x the ONH rim size. This minimizes bias from the relatively intense ONH signal, which can extend slightly outside the segmented edge of the ONH. In the macular region, GAF intensity was calculated as the average GAF intensity over a 4.8 mm diameter region centered around the fovea. Additionally, the macular stress index was calculated and used as an indicator of fluorescence heterogeneity. This proprietary, dimensionless index reflects the spatial variability of GAF intensity within the same region, with higher values indicating greater heterogeneity.

### OCT and retinal layer thickness measurement

Blue autofluorescence cSLO imaging was performed prior to OCT, and infrared SD-OCT combined with cSLO was then acquired with the corneal curvature set to 6.5 mm.^13^ Spectralis blue light fundus autofluorescence images (488 nm excitation, emission detected between 570 to 780 nm) were obtained and reviewed to screen for visible retinal autofluorescence abnormalities. Peripapillary OCT imaging used a standard 12° diameter circular B-scan centered on the ONH. Macular OCT imaging used a 30° horizontal raster-scan with 193 B-scans centered on the fovea. Retinal layer thicknesses were manually measured in ImageJ (NIH, Bethesda, MD, USA) at predefined sampling locations and were reviewed by experienced graders as previously described.^13^ Measurements were obtained at four locations on the circumpapillary B-scan (superior, inferior, nasal, and temporal) and at three locations on the horizontal macular B-scan (the foveal center and points located 1.5 mm nasal and temporal to the fovea). Manual segmentation was selected because automated Spectralis segmentation has shown limitations in rhesus macaque eyes; in our prior work, automated and manual Spectralis RNFL measurements differed significantly, with automated segmentation errors observed in healthy macaques as well as a NHP model of autosomal dominant optic atrophy.^13,23^ The layers assessed included the retinal nerve fiber layer (RNFL), ganglion cell layer (GCL), inner plexiform layer (IPL), inner nuclear layer (INL), outer plexiform layer (OPL), outer nuclear layer (ONL), inner segments (IS), outer segments (OS), retinal pigment epithelium (RPE), choriocapillaris (CC), outer choroid (OC), and total retinal thickness (TRT). Composite metrics included the ganglion cell complex (GCC) defined as RNFL+GCL+IPL and the inner + outer segment complex (IS+OS). Anatomical locations used for retinal layer measurements are provided in **Supplementary Figure S1**. Formal intergrader and intragrader reliability metrics were not prospectively collected because this study was designed to establish normative structure-function relationships rather than to validate segmentation repeatability.

To relate regional GAF to retinal structure, we paired each regional GAF intensity with OCT layer thickness measurements from two standardized B-scans: the circumpapillary scan and a horizontal macular scan through the foveal center. At the ONH, the mean of the four circumpapillary values was used for primary comparisons, and the individual locations were included as predictors in multivariate analyses. In the macula, the two parafoveal measurements were averaged for primary analyses, and the nasal, temporal, and foveal center measurements were also entered individually in multivariate models. For the PFB, GAF was compared with the temporal circumpapillary measurement. Because the OcuMet Beacon does not provide precise axial discrimination, these OCT correlations were interpreted as structural associations with the integrated regional GAF signal rather than as evidence that the GAF originated from a specific retinal layer or cell population.

### Functional testing by PERG and full field ERG with PhNR

Pattern ERG (PERG) and photopic full field ERG (ffERG) were obtained within three weeks of the GAF session. Both electrophysiology tests were recorded under intramuscular sedation followed by an intravenous ketamine constant rate infusion.

The PERG was recorded under photopic conditions. Monocular pattern-reversal stimulation was delivered with a RETIport/scan 21 Q450 stimulator (Roland Consult Stasche & Finger GmbH, Brandenburg an der Havel, Germany). The field size was 52 degrees, with check sizes of 48 arc minutes and 120 arc minutes. Each run delivered 20 seconds of stimulation, with 36 scan lines per square. Peak times for N35, P50, and N95 were recorded, and amplitudes were measured from N35 to P50 and from P50 to N95.

Photopic full-field ERG was performed using the RetEvet handheld unit (LKC Technologies, Gaithersburg, MD). The macaques underwent a ffERG protocol specifically designed to assess the photopic negative response (PhNR), using multiple light intensities that ranged from 0.001 to 12.56 cd·s/m², with 50 flashes per intensity delivered at 2 Hz. The PhNR was defined as the trough after the B-wave within 55 to 90 ms of flash onset, measured from baseline to assess retinal ganglion cell function. For analysis, PhNR amplitude was extracted at 72 ms and at the minimum following the B-wave.

### Statistical analysis

Descriptive statistics summarized demographic and ocular variables; we report the mean ± standard deviation (SD) and, where appropriate, the median and interquartile range (IQR). Normality was assessed with the Shapiro-Wilk test. Bivariate associations were computed with Spearman rank correlation for nonparametric data and with Pearson correlation when normality was achieved. To further explore potential non-linear patterns, we additionally applied LOESS (locally estimated scatterplot smoothing), a method that fits local regressions across the dataset to highlight changes in trend. For all analyses, linear mixed-effects models (LMMs) were used to account for repeated measures within animals and inter-eye correlation. In analyses with a single observation per animal, generalized estimating equations (GEEs) with clustering by subject were applied to account for inter-eye correlation. A *P* value of less than 0.05 defined statistical significance. Multiplicity was controlled using Bonferroni adjustment for prespecified primary comparisons and Benjamini Hochberg false discovery rate control at 5 percent for families of related exploratory tests; reported *P* values are adjusted *P* values.

We additionally performed a thorough multivariate predictor analysis comparing the GAF in each subregion to all the other functional and structural parameters. All analyses were performed in R version 4.4.1, with figures produced in R and GraphPad Prism version 10.4.0.^24,25^ Data entry and preliminary quality checks were managed in Microsoft Excel 2024.

## RESULTS

### Demographic characteristics, image acquisition and quality control

Eighty-two rhesus macaques (13 males and 69 females) were included in the study (**Table 1**). All age groups were represented: 5 infants (6.1%, <1 year), 16 juveniles (19.5%, 1-5 years), 32 adults (39.0%, >5 to <19 years), and 29 geriatrics (35.4%, ≥19 years). Female macaques predominated in each category, particularly in the adult (n=21) and geriatric (n=27) groups, reflecting the female overrepresentation of a breeding colony. Median age of the cohort was 12.90 (IQR 7.27–18.57, range 0.11–29.39) years; mean ± SD was 12.76 ± 7.18 years.

**Table 1:**
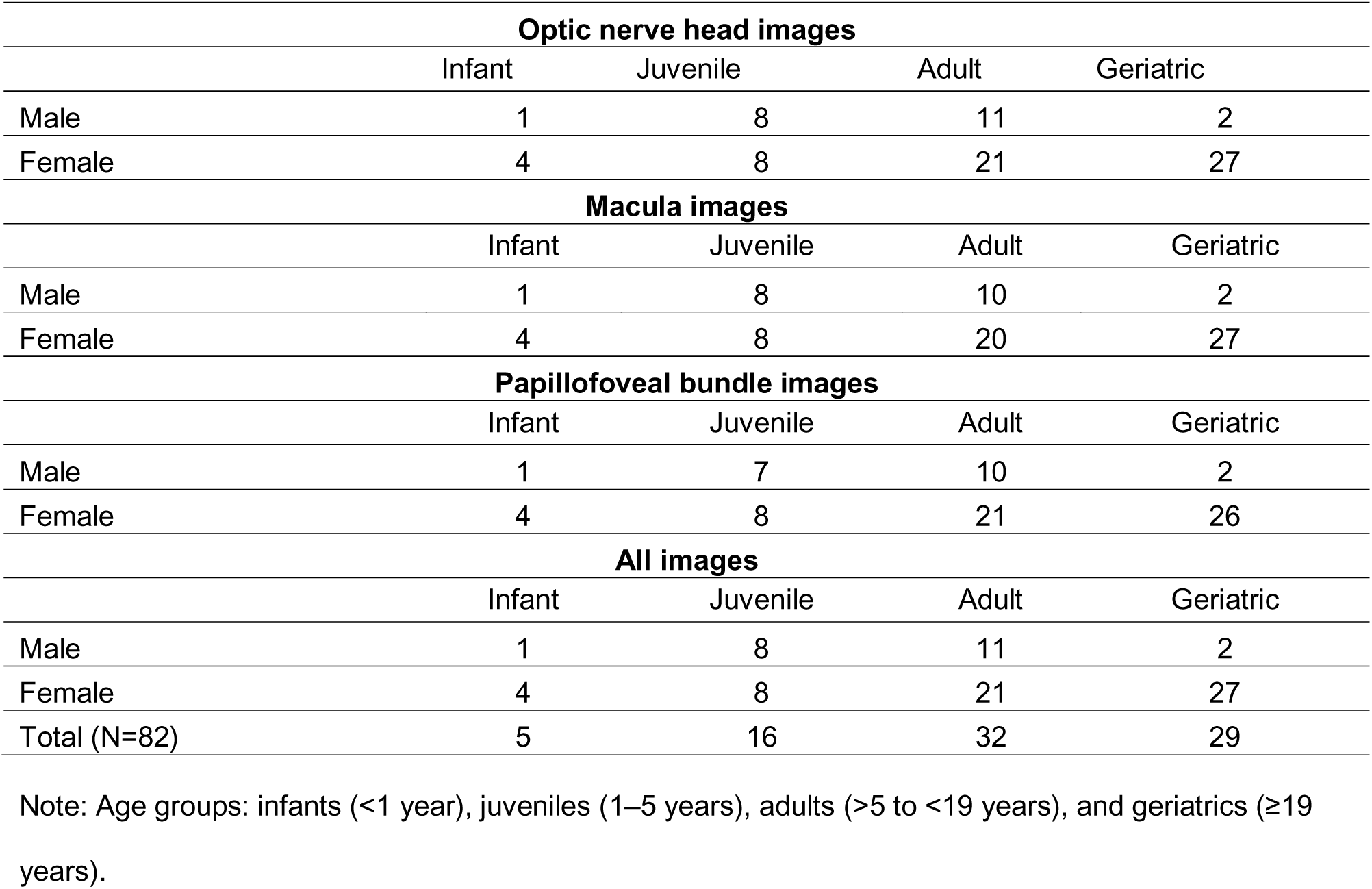
Age groups represented per area imaged.

A total of 209 imaging sessions from 82 rhesus macaques were included after quality review. Of these, 30 rhesus macaques underwent repeated imaging sessions: 2 sessions (n=19), 3 sessions (n=4), 5 sessions (n=2), 6 sessions (n=1), and 7 sessions (n=4). Blue autofluorescence cSLO and OCT images were reviewed to screen for visible retinal or RPE abnormalities, including drusen-like deposits, focal hyperautofluorescent lesions, or structural changes that could suggest irregular lipofuscin accumulation or other retinal pathology. No such clinically apparent abnormalities were identified. This does not exclude physiological background autofluorescence or minor contributions from lipofuscin or other endogenous fluorophores to the measured GAF signal. Imaging acquisition in rhesus macaques was challenging and required multiple attempts in some individuals. Imaging was incomplete in two males: macular acquisition failed in one adult and papillofoveal acquisition failed in one geriatric, in both cases due to an elongated rostrum that prevented the necessary probe angulation. Among images that passed quality control, the mean ± SD GAF image quality indices were 60.66 ± 7.63 for the ONH, 66.37 ± 6.49 for the macula, and 62.95 ± 9.39 for the PFB (**Suppl. Table S1**). When comparing interocular reliability by pairing OD and OS within each subject and computing OD-OS differences, we observed that macular GAF showed a mild OD>OS asymmetry with higher GAF intensity in OD (mean OD-OS +0.86 gsu; *P*=0.038), whereas no significant interocular differences were detected for the ONH (*P*=0.055) or the PFB (*P*=0.35).

Some GAF images were acquired under intramuscular (IM) sedation as required for unrelated studies. However, pronounced eye movements made image acquisition difficult, prompting us to evaluate whether general anesthesia (GA) could facilitate the process. Therefore, we compared two sedation/anesthesia protocols: (1) IM sedation with midazolam (0.1 mg/kg), ketamine (5–30 mg/kg), and dexmedetomidine (0.015 mg/kg), and (2) IM sedation followed by maintenance with isoflurane via endotracheal intubation. Comparing both, acquisition was consistently more challenging under IM sedation because of persistent ocular movements. Nevertheless, all the images were successfully acquired and in those that passed quality control, GAF intensity outcomes did not differ significantly between the two protocols (**Suppl. Table S2**). Accordingly, imaging was performed under GA whenever feasible to minimize eye movement; when GA was not possible, sessions under IM sedation were obtained, and data from both protocols were combined for subsequent analyses.

### Age and GAF are positively correlated

For reference, representative images of all three regions are shown in **Fig. 1**. The ONH GAF intensity showed a significant positive association with age (R=0.39, *P*=2.44×10^-4^; **Fig. 2A1**). The LOESS curve closely paralleled a linear fit, meaning that each additional year of age corresponds to an approximately constant increase in ONH GAF (**Fig. 2A2**). In the macula, GAF intensity showed a strong age-related effect (R=0.63, *P*<1×10⁻¹□; **Fig. 2B1**). When we applied the LOESS regression (**Fig. 2B2**), macular GAF rose steeply during development and early adulthood (0–15 years), plateaued in mid-adulthood (15-19 years), and declined slightly in geriatric rhesus macaques (≥19 years). In the PFB, GAF intensity also increased significantly with age (R=0.66, *P*<1×10⁻¹□; **Fig. 2C1**). The LOESS regression (**Fig. 2C2**) demonstrated that PFB GAF rose steeply during early life and juvenile-adults (∼0–15 years), plateaued in mid adulthood (∼15-19 years), then decreased slightly in geriatric rhesus macaques (≥19 years).

**Figure 1.**
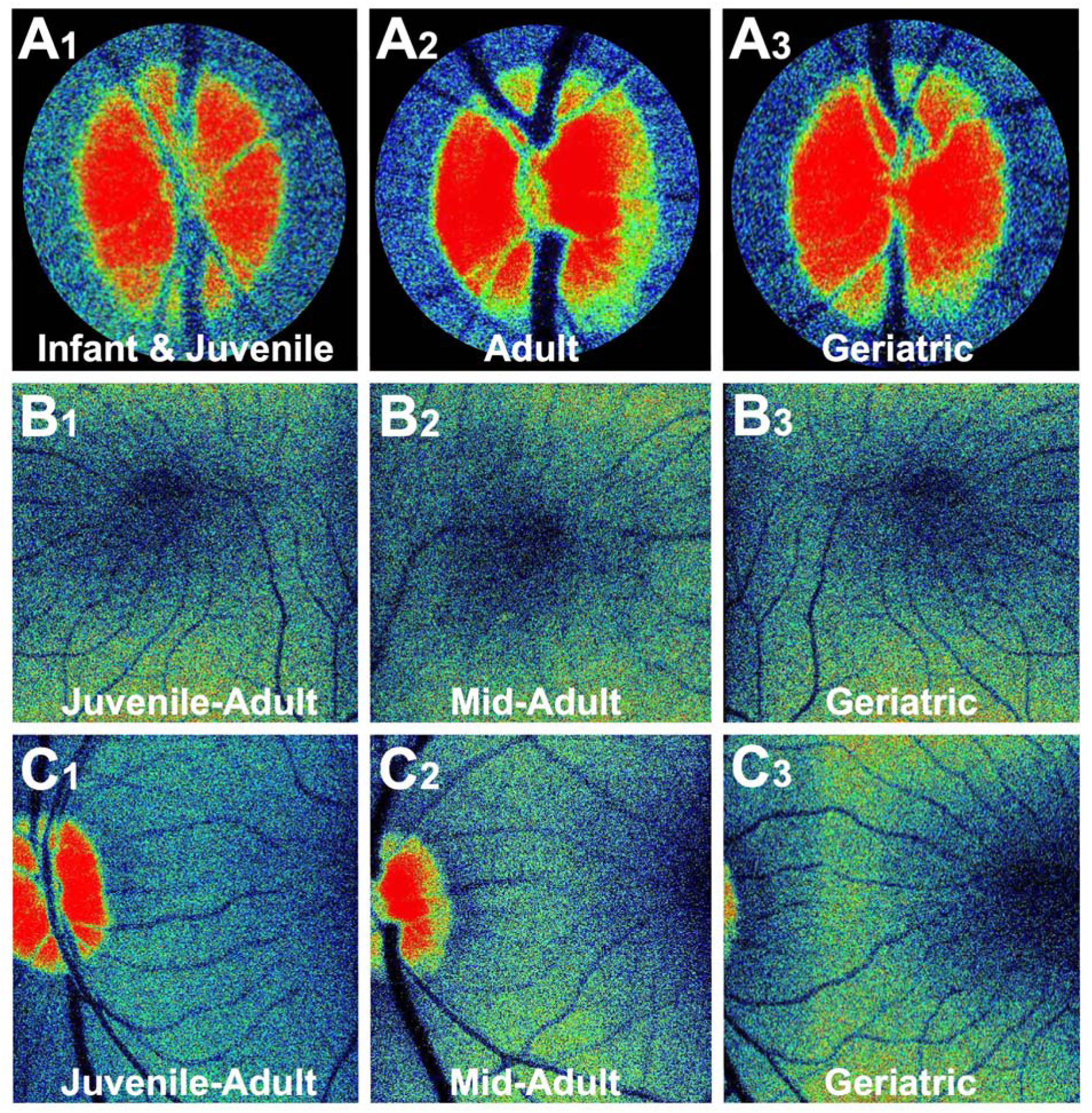
Representative regions of interest for green autofluorescence (GAF) across age groups. **A1-A3**, optic nerve head (ONH); **B1-B3**, macula; **C1-C3**, papillofoveal bundle (PFB). Each panel shows a representative raw GAF image from a rhesus macaque in the different life stages. Age groups: infants & juveniles (<1-5 years), juvenile-adults (0-15 years), mid-adults (>15 to <19 years), adults (>5 to <19 years), and geriatrics (≥19 years). The analysis algorithm automatically detects and quantifies the predefined region of interest even when the acquisition is not perfectly centered, ensuring consistent sampling across eyes and sessions.

**Figure 2.**
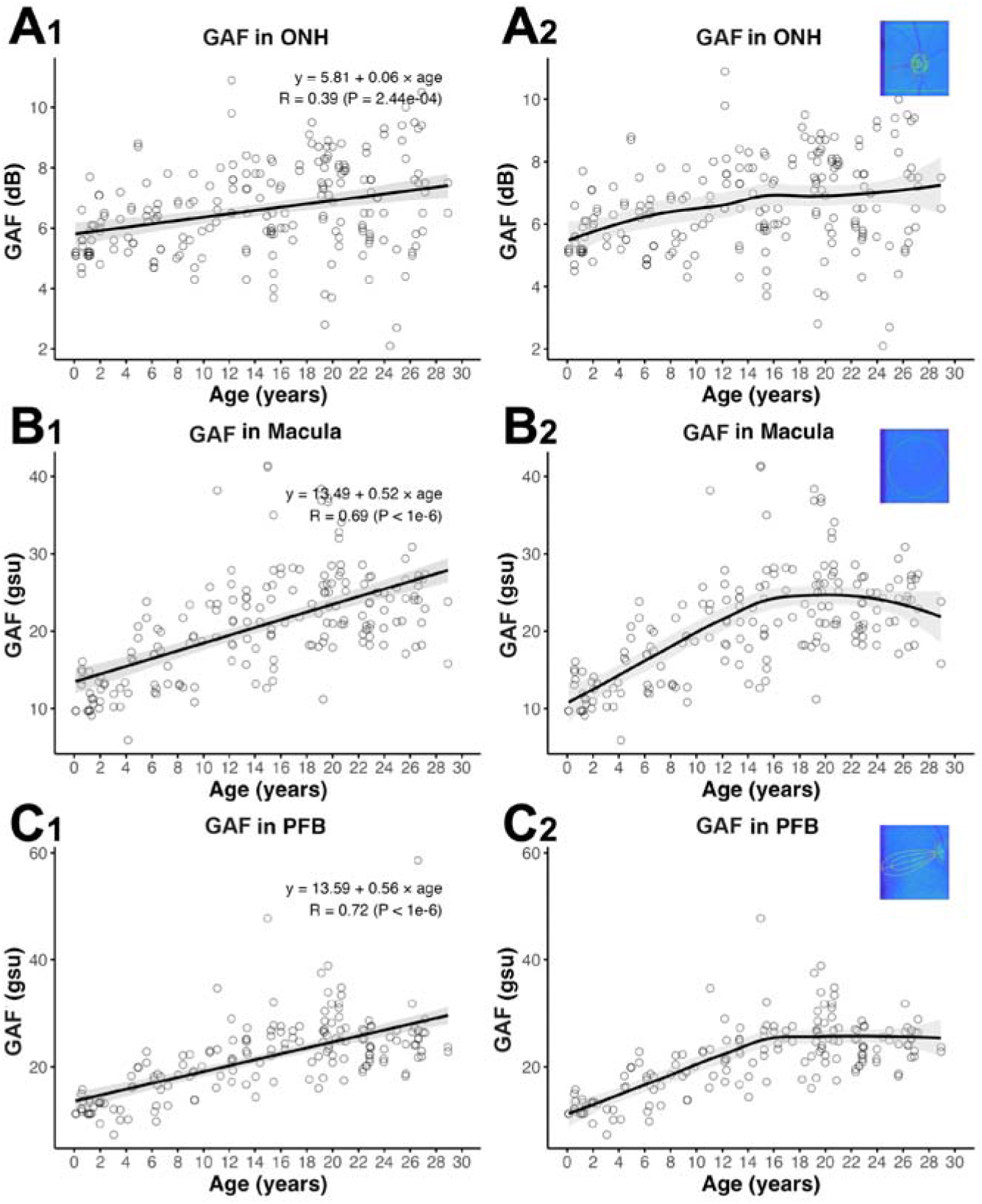
Age influences green autofluorescence (GAF) in normal rhesus macaques, increasing nearly linearly at the optic nerve head (ONH), but in the macula and papillofoveal bundle (PFB) it rises, plateaus in mid adulthood, and declines in geriatrics. Panels **A1** and **A2** present GAF measurements from the ONH (decibels, dB) as a function of age; panels **B1** and **B2** present corresponding data from the macula (grayscale units, gsu) and panels **C1** and **C2** correspond to the PFB (gsu). In each panel, open circles indicate individual eyes imaged. Panels **A1, B1** and **C1** overlay linear mixed-effect models (LMMs) regression lines with shaded 95 % confidence bands; panels **A2, B2** and **C2** display LOESS fits with their confidence bands. LMMs models were clustered by animal to account for correlation between fellow eyes and repeated measurements within the same individual. The regression formula of the LMM-adjusted linear regression and the correlation coefficient (R) with associated *P*-value are annotated in each panel (**A1-B1-C1**). Insets show representative pseudocolor GAF images from the analyzed regions for reference.

For visualization of individual trajectories over time, we included only rhesus macaques with at least three imaging sessions (n=11). Although interindividual variability was evident, within eye longitudinal trajectories were stable or rose slightly with age in the ONH (**Suppl. Fig. S2A**), macula (**Suppl. Fig. S2B**), and PFB (**Suppl. Fig. S2C**), with modest within eye fluctuations. Overall, these trajectories agree with the positive age gradients estimated in the cross-sectional analysis and are most apparent in the macula and the PFB.

We then assessed whether ocular growth influences the association between macular GAF and age in the fixed macular scan by applying an axial length-based magnification correction derived from published rhesus biometry and an independent colony dataset (**Supplementary Data S1**); the age association remained significant but was only modestly attenuated after correction (raw slope=+0.52 gsu/year; corrected slope=+0.453 gsu/year), although future studies should obtain axial length at the same GAF imaging session to enable subject specific correction.

### Sex did not influence GAF across regions

Since our population was weighted towards proportionally more older females, to explore whether GAF varied by sex, we performed an age (±6 months) and sex-matched cohort comparison. In the ONH, mean GAF values were 5.72 dB in males (n=17) and 6.36 dB in females (n=14) and did not significantly differ (*P*=0.1484, **Fig. 3A**). In the macula, mean values were 16.41 gsu in males (n=17) and 18.31 gsu in females (n=13), also not significantly different (*P*=0.4497, **Fig. 3B**). In the papillofoveal bundle (PFB), mean values were 17.59 gsu in males (n=14) and 17.79 gsu in females (n=13), again showing no significant difference (*P*=0.9346, **Fig. 3C**). Since no sex effect was identified across any region, data from both sexes were combined for all subsequent analyses.

**Figure 3.**
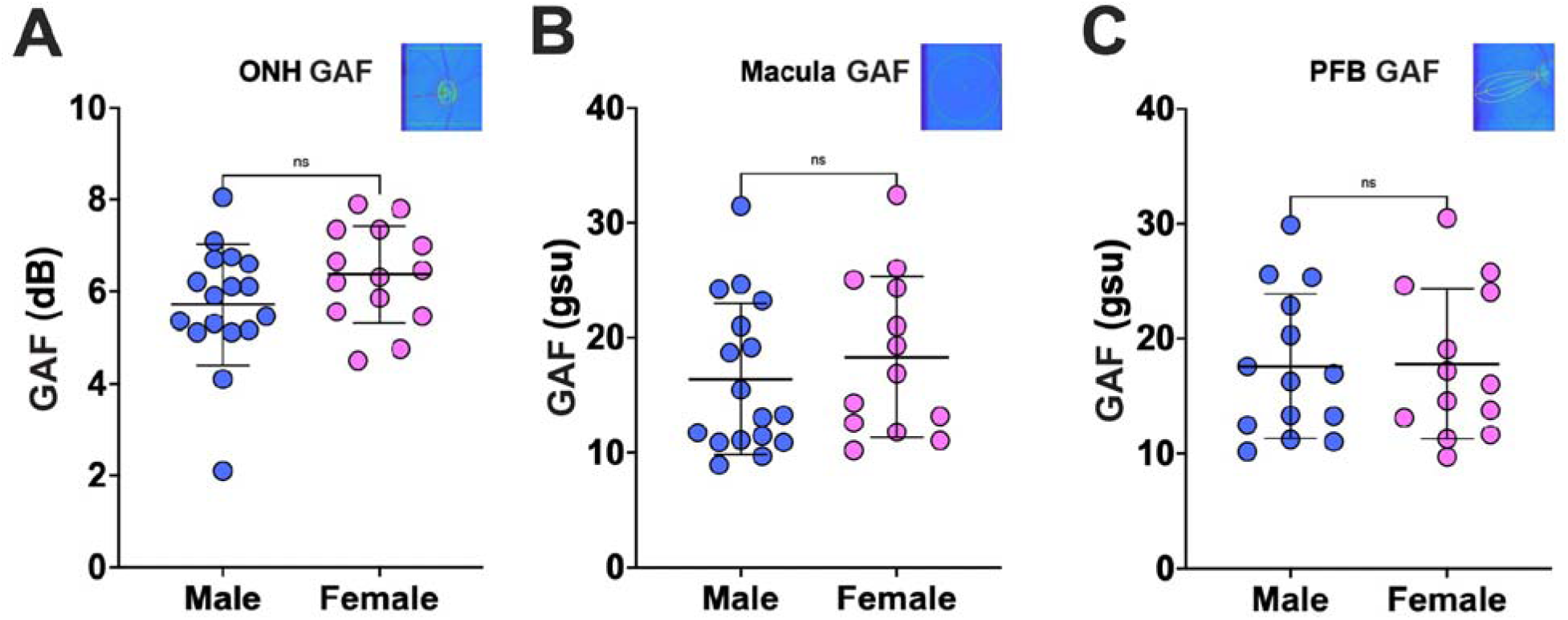
After analyzing the data by age and sex matched cohorts, green autofluorescence (GAF) intensity did not differ between sexes at the optic nerve head (ONH), macula, or papillofoveal bundle (PFB). **A**, ONH GAF (dB); **B**, macular GAF (gsu) intensity; and **C**, papillofoveal bundle (PFB) GAF (gsu). Data are shown as individual values with mean ± SD. Insets show representative pseudocolor GAF images from the analyzed regions for reference.

### Regional associations between GAF and retinal layer thickness measured by OCT

We related regional GAF to retinal structure by comparing GAF intensity with OCT layer thickness measured at predefined locations. All correlations are shown in **Suppl Fig. S3** (ONH), **Suppl Fig. S4** (macula), and **Suppl Fig. S5** (PFB). After adjusting for age, global ONH GAF generally increased as neuroretinal layers in the circumpapillary scan became thinner (**Suppl. Fig. S3**). Among photoreceptor metrics, OS reached significance (**Fig. 4A1**; **Suppl. Fig. S3I**; *P*=0.025), and the combined IS+OS metric showed the strongest effect (**Fig. 4A2**; **Suppl. Fig. S3J**; *P*=0.005). Total retinal thickness was also negatively correlated with GAF (**Fig. 4A3**; **Suppl. Fig. S3L**; *P*=0.008). In the macular region, GAF showed predominantly negative associations in the outer retina, with small positive trends in selected inner layers (**Suppl. Fig. S4**). The INL was the only layer with a significant positive association (**Fig. 4B1**; **Suppl. Fig. S4E**; *P*=0.025). Significant inverse associations were observed for the OPL (**Fig. 4B2; Suppl. Fig. S4F**; *P*=0.011) and the IS+OS complex, which showed the largest effect (**Fig. 4B3**; **Suppl. Fig. S4J**; *P*<0.001). Lastly, global PFB GAF showed no statistically significant associations with any temporal circumpapillary retinal layer (**Suppl. Fig. S5**), indicating no detectable structure-function relationship in this region.

**Figure 4.**
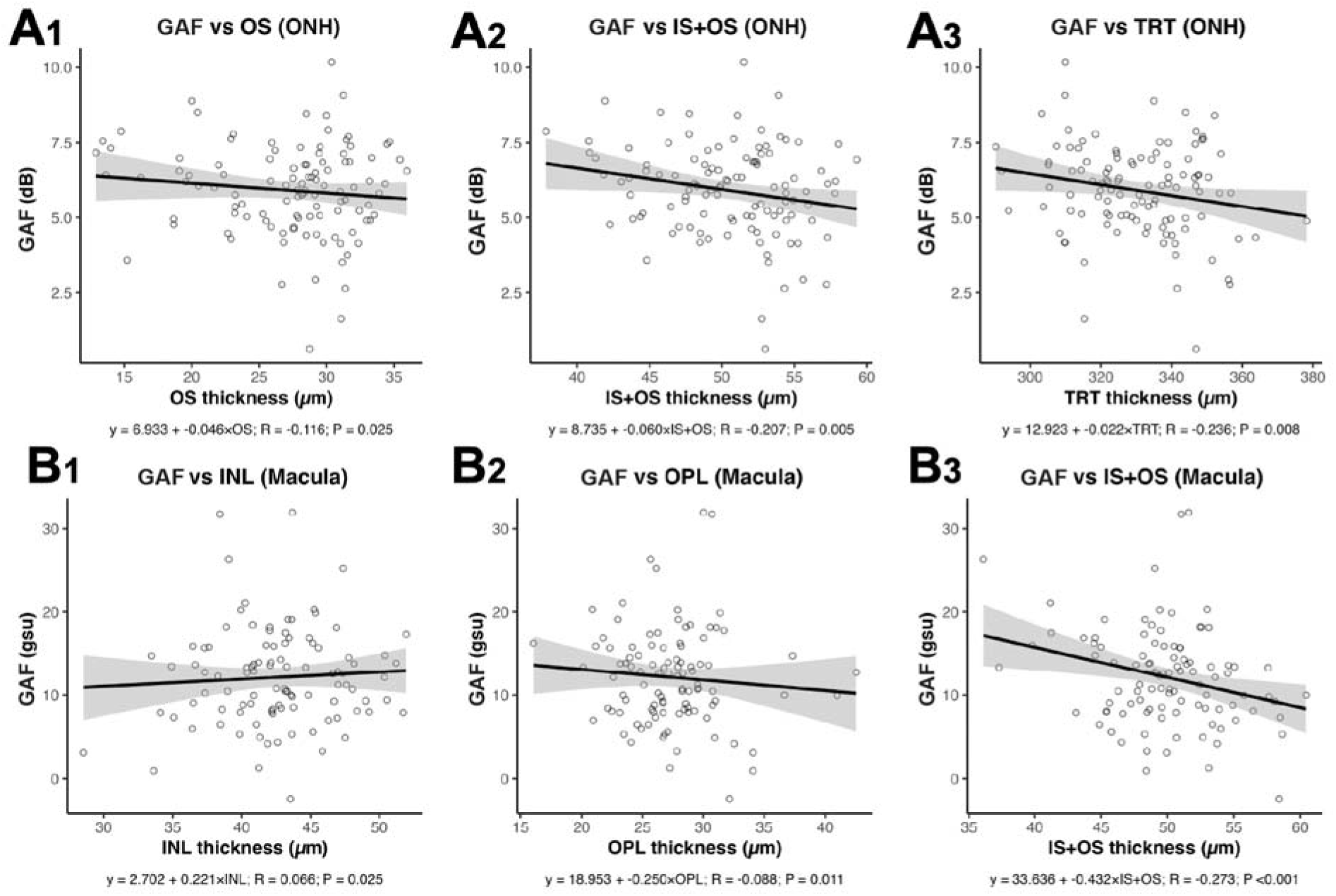
Green autofluorescence (GAF) is inversely associated with outer retinal thickness in both the ONH and macula, while the macular inner nuclear layer (INL) shows a small positive association. **A1-A3**, ON GAF (dB) compared to outer segment (OS) thickness (**A1)**, inner + outer segment (IS+OS) thickness (**A2**), and total retinal thickness (TRT; **A3**). The layer thickness to compare with the ONH GAF was taken from the circumpapillary scan as the mean of four locations: superior, inferior, temporal, and nasal. **B1-B3**, macular GAF (grey scale units, gsu) compared to INL thickness (**B1**), outer plexiform layer (OPL) thickness (**B2**), and IS+OS thickness (**B3**). Retinal layer thickness was measured on the macular OCT scan at two parafoveal sites located 1.5 mm nasal and temporal to the foveal center; the mean of these two measurements was used for analysis. Each point represents one eye at a single visit in which OCT and macular GAF were obtained. Solid lines show linear fits from generalized estimating equations that account intereye correlations within subjects, with 95% confidence bands. Panel annotations report the model regression formula, together with the R and *P*-values.

### Association of IOP with GAF after age adjustment

After age adjustment, GAF did not correlate with IOP at the ONH and PFB (*P*=0.8404, **Fig. 5A** and *P*=0.1721, **Fig. 5C**, respectively). By contrast, in the macula, higher IOP was significantly associated with higher GAF (*P*=0.0230; **Fig. 5B**). Across regions there was a clear tendency for higher IOP with higher GAF; however, inference was limited because only three macaques showed IOP values above the physiological range (interocular mean 23, 23.5 and 28 mmHg), and these elevations were transient. The mean IOP in the whole cohort was 14.8 ± 3.54 mmHg (range 6-32 mmHg). For context, normal IOP in rhesus macaques is 15-16 mmHg, placing the upper end of normal at 22 mmHg.^18,26^ We repeated the correlation analysis after excluding IOP measurements above the upper end of normal (**Suppl. Fig. S6**) and found that the ONH association remained nonsignificant (**Suppl. Fig. S6A**), whereas positive associations were retained in the macula (slope=+0.28 gsu/mmHg, R=+0.30, *P*=0.0147; **Suppl. Fig. S6B**) and were also detected in the PFB (slope=+0.24 gsu/mmHg, R=+0.18, *P*=0.0400; **Suppl. Fig. S6C**).

**Figure 5.**
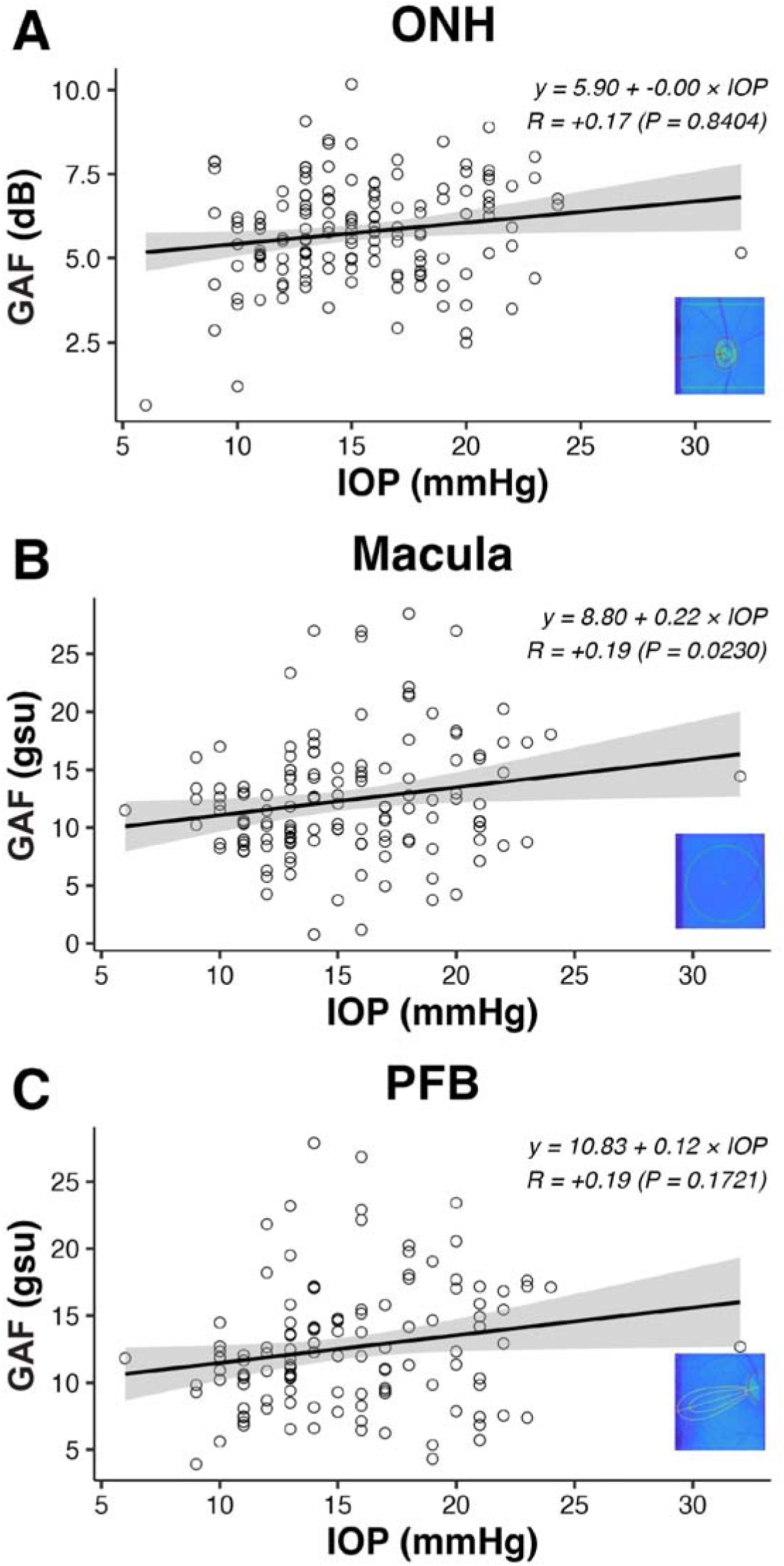
After adjusting for age, green autofluorescence (GAF) shows significant association with intraocular pressure (IOP) in the macula. **A**, optic nerve head (ONH, dB); **B**, macula (grey scale units, gsu); **C**, papillofoveal bundle (PFB, gsu). Each data point represents one eye at a single imaging session with IOP recorded at the same visit. Solid lines show linear fits from mixed model accounting for intereye and intraindividual repeated measures correlation with 95 percent confidence bands. Insets show representative pseudocolor GAF images from the analyzed regions for reference.

### Electrophysiologic correlates of GAF by region

We evaluated the relationship of GAF intensity with PERG in 19 rhesus macaques. Across regions, only the ONH showed statistically significant associations. The ONH GAF was inversely related to the N35-P50 (**Figure 6A1**; R=−0.33; *P*=0.0294) P50-N95 (**Figure 6A2**; R=−0.47; *P*=0.0239) components consistent with higher GAF associating with lower PERG amplitudes. No significant associations were identified between GAF intensity and the PhNR (n=19) (**Suppl. Fig S7**).

**Figure 6.**
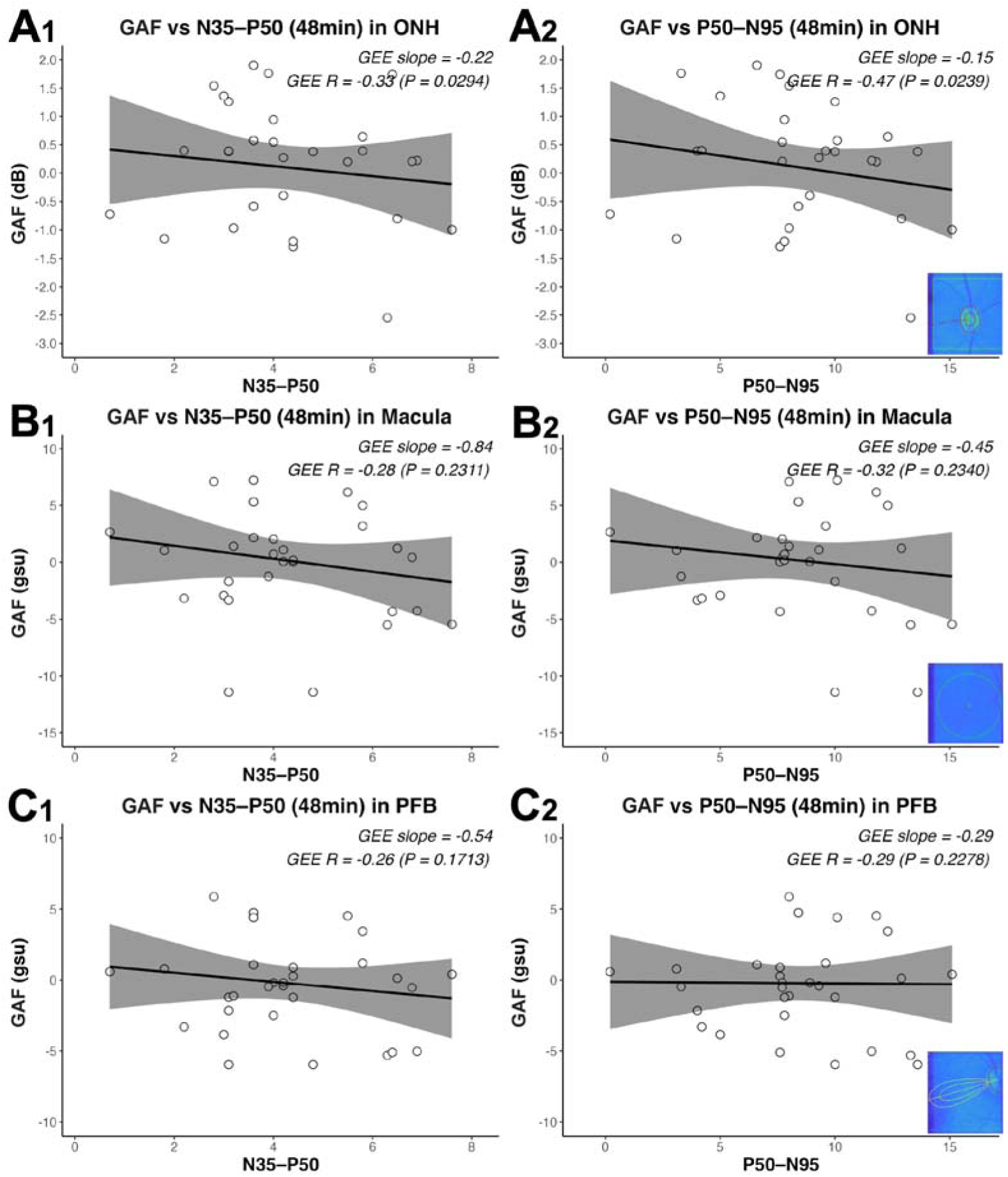
Higher green autofluorescence (GAF) in the optic nerve head (ONH) is associated with lower pattern ERG (PERG) amplitudes. **A1**, optic nerve head (ONH) GAF (dB) versus PERG N35–P50 amplitude (µV); **A2**, ONH GAF (dB) versus PERG P50–N95 amplitude; **B1**, macula GAF (gsu) versus PERG N35–P50 amplitude; **B2**, macula GAF (gsu) versus PERG P50–N95 amplitude; **C1**, papillofoveal bundle (PFB) GAF (gsu) versus PERG N35–P50 amplitude; **C2**, PFB GAF (gsu) versus PERG P50–N95 amplitude. Each data point represents one eye at a single imaging session; PERG testing and GAF imaging were performed within a maximum interval of three weeks. Solid lines show linear fits from linear mixed models (LMM) accounting for intereye correlation with 95 percent confidence bands. Annotations in each panel report the LMM regression (GAF units per µV), marginal R, and *P-*values. Insets show representative pseudocolor GAF images from the analyzed regions for reference.

### Normative green autofluorescence by age group

Using the standardized workflow to center, image, and quantify GAF intensity in the ONH rim (**Suppl. Fig. S8A-C**), circumferential profiles showed a stable spatial topology across age groups: a temporal peak, an intermediate nasal, and minimal in the superior and inferior sectors (**Suppl. Fig. S8D**). For context, in a human cohort processed with the same workflow, quadrant means were 8.0 dB in the temporal, 6.8 dB in the nasal, 5.1 dB in the inferior, and 5.0 dB in the superior (n=69; unpublished data, OcuSciences). Quantitative quadrant comparisons confirmed this pattern in every age group (**Suppl. Table S3** and **Fig. 7**). In infants and juveniles, temporal significantly exceeded superior and inferior (**Fig. 7A1**; *P*<0.01; normative values detailed in **Suppl. Table S3**), consistent with the reference map in **Fig. 7A2**. Adults showed the same ordering with larger separations (**Fig. 7B1-B2**; **Suppl. Table S3**). Geriatric macaques also maintained this pattern (**Fig. 7C1-C2**; *P*<0.05; **Suppl. Table S3**). In our adult cohort, the global mean was 6.51 ± 1.31 dB; by comparison, a human cohort analyzed with the same algorithms showed 4.4 dB.

**Figure 7.**
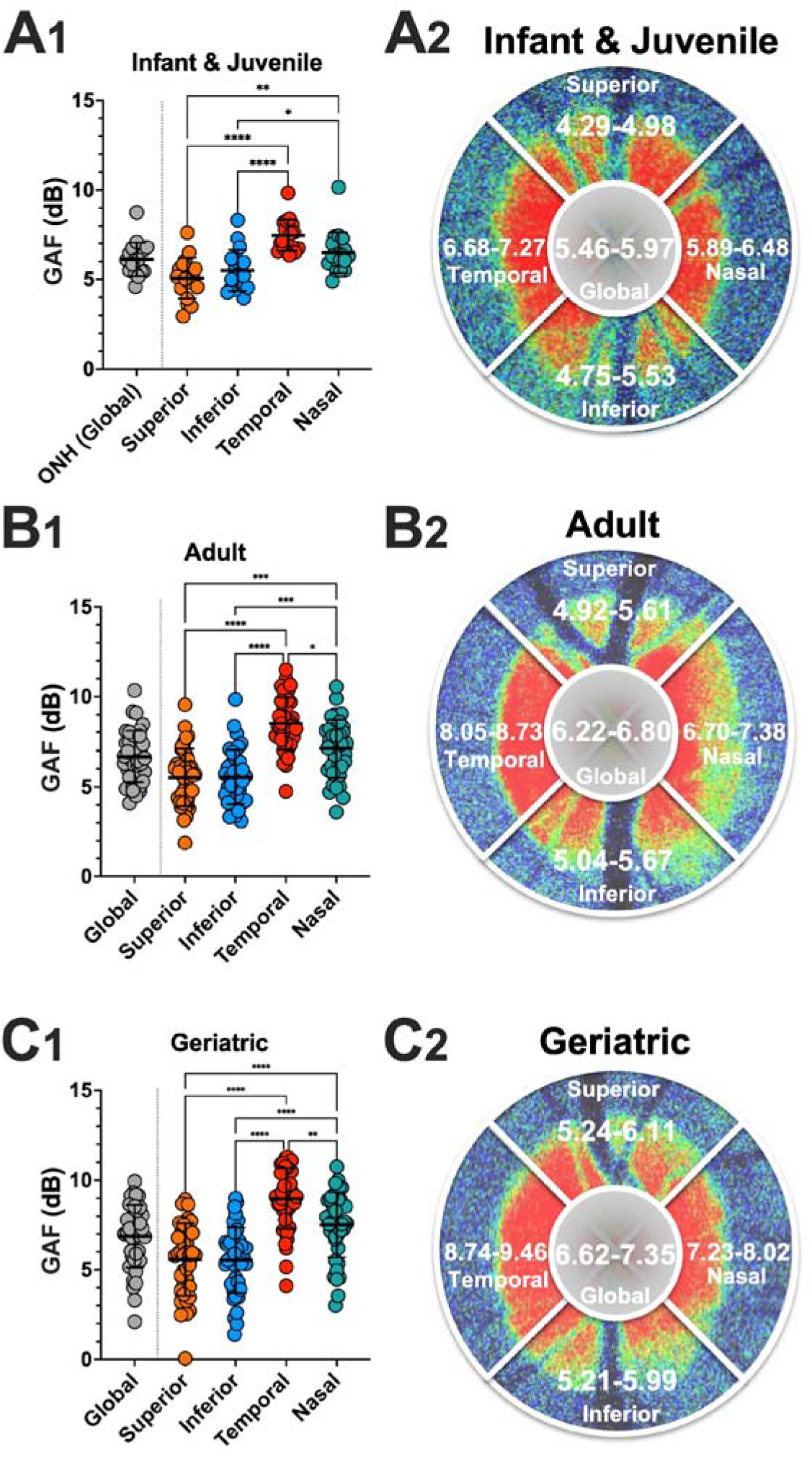
Normative green autofluorescence (GAF) in the optic nerve head (ONH) varies by quadrant across all age groups, with the temporal quadrant consistently highest. **A1**, regional variation within Infant and Juvenile across Superior, Inferior, Temporal, and Nasal; the global value is the average of these four sectors. **A2**, reference map with sector and global values shown with the central 95 percent reference interval over a representative GAF image for Infant and Juvenile. **B1** and **B2**, same for Adult. **C1** and **C2**, same for Geriatric. In **A1**, **B1**, and **C1**, each point represents one eye at one imaging session; the thick line represents the mean and the error bars represent the standard deviation; *P* values (* less than 0.05, ** less than 0.01, *** less than 0.001) are from mixed models accounting for intereye correlation and intraindividual repeated measures. Age groups: infants (<1 year), juveniles (1–5 years), adults (>5 to <19 years), and geriatrics (≥19 years).

As expected, across age groups, macular GAF intensity increased with age, juvenile adults show the lowest values, with higher means in mid adults and geriatrics (**Suppl. Table S4, Fig. 8A1-C1**), and the central 95 percent reference intervals shift upward accordingly (**Fig. 8A2-C2**). By contrast, macular GAF heterogeneity shows broad overlap across groups, indicating no clear age dependence in this cohort (**Fig. 8A1-C1**).

**Figure 8.**
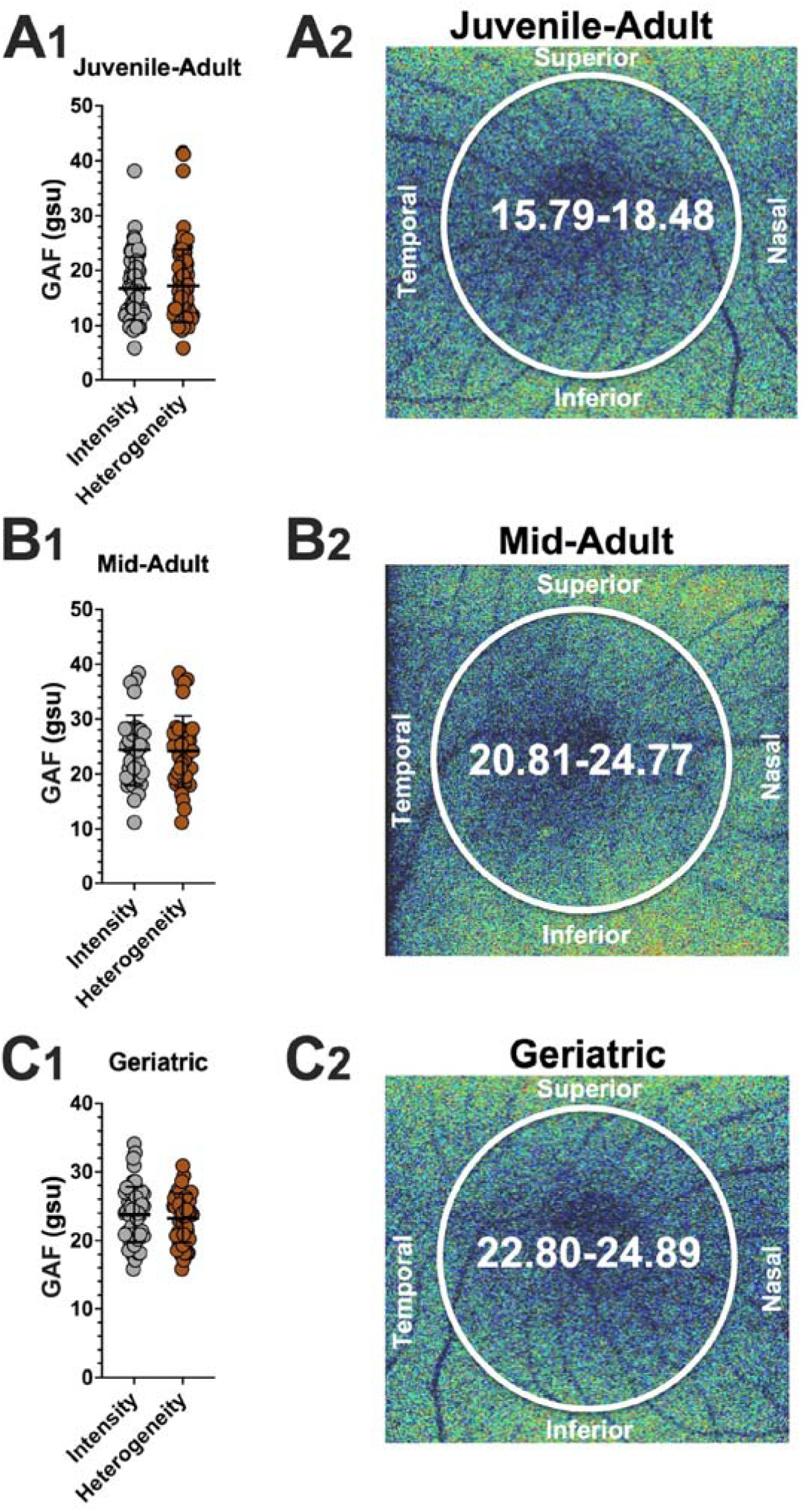
Normative macular green autofluorescence (GAF) varies with age, being higher in mid-adult and geriatric groups than in juvenile-adults. **A1**, regional summary of two metrics within the circular region of interest: GAF intensity (mean GAF within the region) and GAF heterogeneity (spatial variability of GAF within the same region). **A2**, central 95 percent reference interval for GAF intensity over a representative raw GAF image. **B1** and **B2**, same for Mid adult. **C1** and **C2**, same for Geriatric. In **A1**, **B1**, and **C1**, each point represents one eye at one imaging session; the thick line represents the mean and the error bars represent the standard deviation. Age groups: juvenile-adults (0-15 years), mid-adults (>15 to <19 years), and geriatrics (≥19 years).

Similarly, global PFB GAF intensity increased with age; juvenile adults had lower values than mid adults and geriatrics (**Fig. 9A1-C1**; **Suppl. Table S5**). Sector means followed the same pattern of increase with age, with horizontal bundles tending to be higher than arcuate bundles within each age group, although these sector differences did not reach statistical significance (**Fig. 9A1-C1**). The reference maps displayed the central 95 percent intervals for each sector (**Fig. 9A2-C2**).

**Figure 9.**
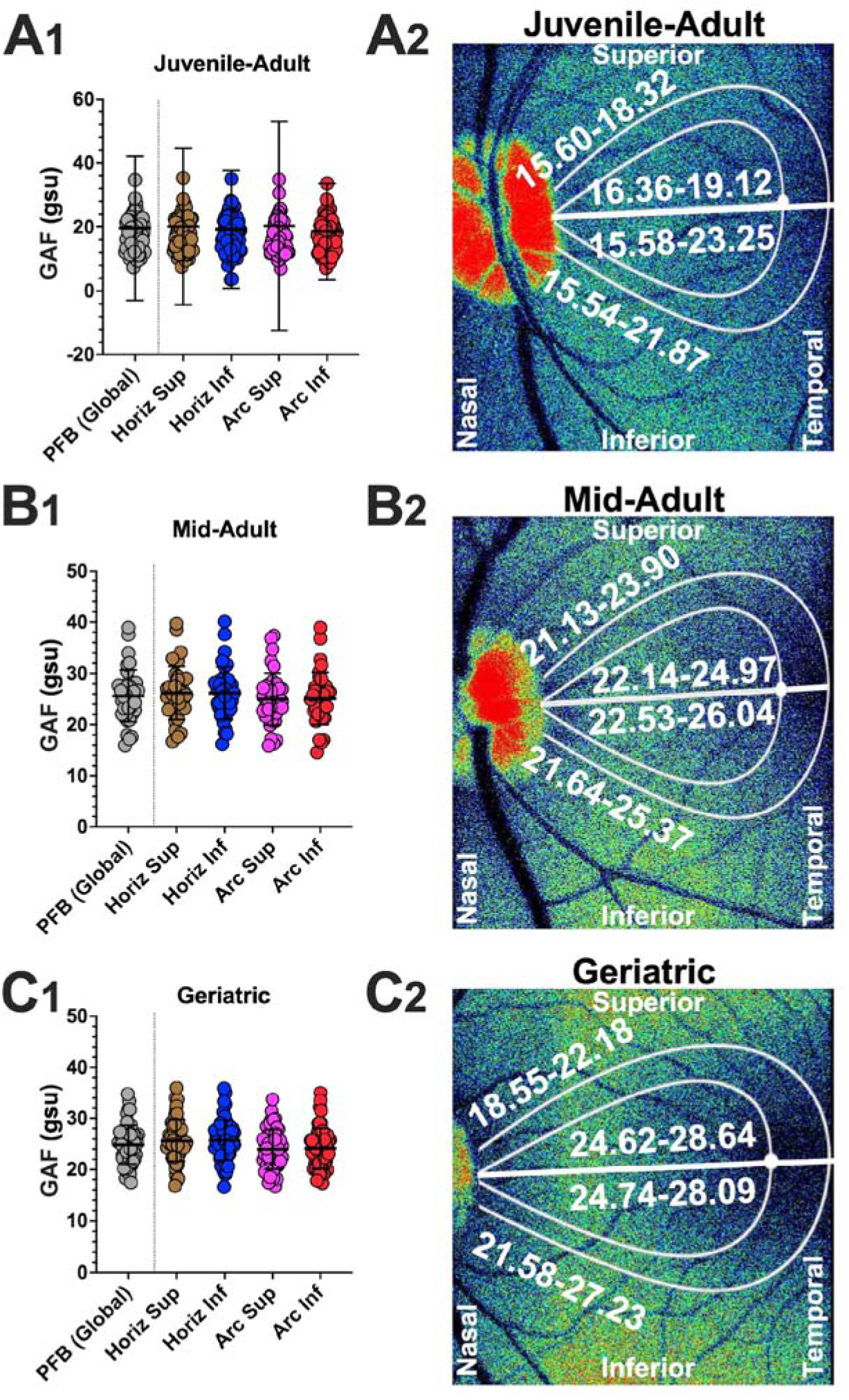
Normative green autofluorescence (GAF) in the papillofoveal bundle (PFB) varies by sector and increases with age. **A1**, regional summary across PFB global and the four predefined subregions Horizontal Superior, Horizontal Inferior, Arcuate Superior, and Arcuate Inferior; the global value is the average of the four subregions and is separated from them by a vertical dotted line. **A2**, reference map with the same masks showing the central 95 percent reference interval for each region over a representative raw GAF image. **B1** and **B2**, same for Mid Adult. **C1** and **C2**, same for Geriatric. In **A1**, **B1**, and **C1**, each point represents one eye at one imaging session; the thick line represents the mean and the error bars represent the standard deviation; no significant differences were found. Age groups: juvenile-adults (0-15 years), mid-adults (>15 to <19 years), and geriatrics (≥19 years).

### Predictors of Green Autofluorescence Across Regions

We evaluated associations between regional GAF outcomes and a comprehensive set of structural and functional parameters using univariate linear mixed effects models that accounted for intereye correlation and repeated sessions, with all models adjusted for age. In total, 264 predictors to outcome combinations reached statistical significance in the age adjusted analyses at the ONH (**Suppl. Table S6**), macula (**Suppl. Table S7**) and PFB (**Suppl. Table S8**).

Across optic ONH sectors, GAF intensity was most strongly linked to retinal thickness measures. Thinner RNFL generally corresponded to higher GAF. Shorter full field ERG A-wave implicit time corresponded to higher GAF in several ONH sectors. For macular GAF heterogeneity, the strongest association was a negative relationship with IOP. Heterogeneity also increased with thicker circumpapillary RPE. It also rose with thicker inner retinal layers on the circumpapillary scan. At the PFB, GAF intensity showed its strongest and most consistent inverse relationships with outer retinal structure measured at the macula. Shorter parafoveal OS corresponded to higher GAF intensity in the PFB. Thicker RPE at the temporal macula and at the parafoveal mean showed positive associations across sectors and globally. Higher IOP was positively associated with GAF intensity in several sectors but not in the global outcome of the PFB. Electrophysiologic measures were directionally concordant, longer full field ERG A- and B-wave implicit time corresponded to higher PFB intensity in some sectors.

## DISCUSSION

Across 82 normal rhesus macaques, GAF increased with age in all regions. After age adjustment, GAF showed a coherent cross regional pattern of association with ocular structure and function. Higher GAF was associated with thicker INL and RPE, and with thinner IS+OS and TRT. Intraocular pressure was positively associated with GAF mainly in the macula, and ONH GAF was inversely associated with PERG amplitudes. Taken together, these findings indicate that GAF reflects region specific variation linked to age, retinal structure, intraocular pressure, and inner retinal function, and that these factors should be considered when interpreting the signal.

Contrary to reports in human patients,^7^ image acquisition in rhesus macaques was technically demanding not only because ocular movements degraded targeting and focus but also because craniofacial anatomy with an elongated rostrum complicated alignment, increased working distance, occasionally obstructed the optical path, and predisposed to shadowing. Rigorous exclusion of scans with targeting errors, poor focus, segmentation or registration artifacts, inadequate dilation, or exogenous fluorescence, together with systematic review of blue autofluorescence images that showed no retinal autofluorescence, minimized the risk that lipofuscin or other fluorophores confounded the measurements. Transparent quality thresholds improved reproducibility, reduced the risk of artifactual associations, and provided a benchmark for future studies as image quality metrics become further standardized.

In healthy rhesus macaques, GAF showed clear age dependence that differed by region. Our regional aging curves are broadly consistent with what is known about GAF while extending it in important ways. In the ONH, GAF has been validated mainly in disease cohorts rather than normative aging, leaving age-only trajectories underreported; an evidence gap that our observed linear increase helps fill.^10^ Prior human work showed that macular retinal GAF in clinically normal controls increases with age, supporting our observation that age is a primary determinant of GAF changes.^6^ Whether the rise, plateau, and decline pattern we observe in healthy rhesus macaques also occurs in humans remains unknown. Published GAF protocols typically center the region of interest on the macula or ONH rather than PFB, so no data is available for comparison in this area. Finally, while direct age-normative GAF data in animals are lacking, mechanistic aging signals align with our trends: multiple species, including non-human primates, show age-related retinal and choroidal iron accumulation that promotes ROS generation, a substrate expected to elevate FAD-derived GAF with age.^27,28^ These observations justify region specific life stage groupings for normative presentation and underscore the need for strict age adjustment when interpreting GAF.

In our age-matched rhesus cohort, we found no evidence that sex influenced GAF at the ONH, macula, or PFB. This pattern was consistent with the human GAF literature, where key clinical studies do not report meaningful sex stratified differences in healthy eyes or common disease cohorts.^7,10,29^ However, sex may become important in neurodegenerating retina, as recent experimental work showed that circulating female hormones exacerbated retinal neurodegeneration, underscoring the need to consider endocrine status when interpreting metabolic imaging signals.^30^ Biologically, modest sex effects in normal retinas remain plausible, as sex hormones modulate retinal antioxidant defenses and neuronal resilience, women have been reported to carry more retinal iron across ages; potentially increasing oxidative stress.^31^ Further research comparing specific hormonal levels and metabolic stress measured by GAF could help elucidate this influence.

The coupling between GAF and retinal layer thickness was dominated by two patterns. We found that macaques with a thicker INL showed higher GAF. This co-occurrence is consistent with physiological activity in which Müller cell potassium and water handling produces transient inner nuclear layer expansion,^32,33^ together with increased inner retinal oxygen consumption during visual stimulation.^34,35^ Second, higher GAF was consistently associated with shorter IS+OS likely reflecting light-exposure physiology at imaging, as photopigment bleaching shortens the OS transiently and thickens the RPE.^36–39^ These INL and outer retinal associations most likely reflect a normal rise in metabolic state due to visual stimulation and photobleaching rather than pathology.^40^ This visual stimulation could come from the ambient room light with dilated pupils under anesthesia, BAF cSLO imaging or the flash from anterior segment and fundus photos rather than from the Beacon device. Additionally, the observation of higher macular GAF in OD (examined first) supports an order-dependent photobleaching effect whereby OS imaged later had more time to recover. The inverse relationship of TRT and GAF in the ONH has been also reported in healthy adult patients, although their cohort was not age-corrected and they concluded that the associations might have been mediated by age.^40^ Thus, these findings support GAF as a physiologically informative imaging signal associated with retinal structure, while also indicating that prior light exposure and photopigment bleaching may influence measurements in healthy eyes and should be controlled for or modeled in future studies.^8^ Although photopigment bleaching is one possible explanation for the inverse association between GAF and IS+OS thickness, this was not directly tested in the present study, and other biological or technical factors may also contribute. Standardized pre-imaging conditions, including control of recent light exposure or dark adaptation when feasible, may improve consistency in future studies specifically designed to evaluate this variable; however, further studies directly comparing acquisition conditions are needed before specific recommendations can be made. Additionally, the association between thinner RNFL and higher GAF may reflect increased oxidative or mitochondrial stress in retinal ganglion cells and their axons, reduced screening of autofluorescence arising from deeper retinal layers or the RPE, or a combination of these mechanisms, depending on the anatomical region analyzed.

In our cohort, GAF showed region specific associations with IOP. At the ONH, we observed a positive nonsignificant tendency for GAF to rise with higher IOP, consistent with human glaucoma eyes, where ONH-rim GAF is significantly higher in patients than controls.^10,41^ In the macula, our cohort showed that higher GAF intensities were significantly associated with higher IOP values. Notably, ocular hypertensive human eyes display significantly elevated macular GAF even before any detectable RNFL thinning, implying that GAF, likely reflecting mitochondrial dysfunction, can be identified early, before structural damage.^42^ Regarding the PFB our analysis we observed that in some sectors, higher IOP translated into significantly higher GAF, likely indicating that these high-metabolic-demand axons experience some oxidative stress under elevated IOP. Since only a few animals exceeded the physiological IOP range, inference at the extremes is limited; nonetheless, the cross-species literature supports that pressures beyond normal likely amplify retinal metabolic stress captured by GAF and motivates the inclusion of this confounding factor in future analysis.^42,43^

Regional comparisons between GAF and electrophysiology suggest that ONH GAF is a sensitive metabolic correlate of inner-retinal function, as it is negatively associated with PERG amplitude. This suggests that ONH GAF as a localized readout of RGC stress and is consistent with disease data showing that, as mitochondrial dysfunction worsens, GAF correlates with functional loss.^10^ Notably, most prior clinical studies relate GAF to non-ERG functional measures, such as best-corrected visual acuity, contrast sensitivity, and visual field indices rather than to ERG directly.^7,10,44,45^ In this cohort, biological variability, 1-3 week gaps between tests, and different anesthesia likely weakened GAF–ERG correlations.

Several limitations should be acknowledged including that most imaging was performed under general anesthesia, which can introduce variability in measurements, and axial globe length was not measured. Inhalational anesthesia could potentially decrease mitochondrial metabolism and therefore reduce the fluorescence signal from flavoproteins.^46,47^ Consistent with this possibility, GAF values were slightly lower under inhalational anesthesia in our dataset, although the difference was not statistically significant. Together with our strict monitoring of anesthetic depth and systemic parameters, these findings suggest that anesthetic protocol differences were unlikely to materially bias the reported normative values or the structure-function associations. Subject-specific axial globe length was not measured. Because ocular magnification affects the transverse scaling of OCT images, variability in axial length may have altered the true retinal location of measurements obtained 1.5 mm nasal and temporal to the foveal center. Additionally, although retinal layer thickness measurements were reviewed by experienced graders, formal intergrader reliability was not prospectively assessed, which may have introduced measurement variability. Although the Beacon uses a confocal optical design, the depth of field limits axial discrimination; therefore, the measured GAF reflects an integrated signal across a broad axial extent of the retina rather than a single retinal layer.^48^ Because the excitation and emission spectra of oxidized flavoproteins overlap with those of lipofuscin and other endogenous retinal fluorophores, RPE-derived autofluorescence may contribute to the measured signal in the macular and peripapillary regions. Therefore, the measured GAF signal should be interpreted as an integrated regional GAF signal rather than as a fluorophore-specific or layer-specific measurement of flavoprotein fluorescence. Mechanistic validation of the detected signal and confirmation of fluorophore specificity require dedicated experimental studies that were beyond the scope of this work and are best addressed through manufacturer led investigations and future collaborative basic science efforts, with complementary fluorescence lifetime imaging and phasor analysis representing a promising strategy to improve fluorophore specificity in future work.^49,50^

Despite these restrictions, our study provides the first normative baseline for GAF in the rhesus macaque eye and yields several key insights. We found that GAF increases with age in the ONH, macula, and PFB, although the pattern of age-related change was region-specific. We also identified significant associations between GAF and other ocular parameters, linking higher GAF to structural and functional signs of stress, consistent with clinical observations that mitochondrial stress is associated with early glaucomatous damage and visual dysfunction.^29^ These associations were most compelling at the ONH, where higher GAF was associated with thinner retinal layers and lower PERG amplitudes, supporting ONH GAF as a particularly relevant readout of retinal ganglion cell axonal structure and inner-retinal function. Importantly, because RPE-derived autofluorescence is absent within the ONH itself, these ONH findings are less likely to be explained by RPE contribution than associations observed in macular or peripapillary retinal regions. Given that no universal reference standard for GAF exists across studies and that macaque GAF values appear different than those reported in humans, our spatial ONH profiles and age-segmented normative data fill a critical gap. Overall, these data provide a reference framework for interpretation of GAF in future studies of retinal and optic nerve disease. Careful interpretation, however, requires attention to age and other relevant covariates, as well as recognition that the precise molecular contributors to the measured signal were not isolated in the present study.

## ACKNOWLEDGEMENTS

This work was supported by the National Institutes of Health (NIH) grants U24EY029904 (AM, SMT), R01EY033733 (SMT), P30EY012576, and T35OD010956. GY is supported by NIH R01EY032238 and R01EY036504. Additional support for this research came from the NBRI Base Grant from the NIH, Office of the Director, OD011107 and a contract from Stoke Therapeutics Inc. The content is solely the responsibility of the authors and does not necessarily represent the official views of the funding agencies. Coauthors affiliated with OcuSciences Inc contributed only to technical accuracy and transparency regarding the device and image processing methods, and did not have a role in study design, data acquisition, statistical analysis, interpretation of the findings, or the decision to submit the manuscript.

The authors thank the many members of the Comparative Ophthalmology and Vision Sciences Laboratory for their contributions throughout the study. Special thanks to Monica J. Motta and Michelle Ferneding for coordinating procedures and providing technical assistance. We are grateful to Chrisoula A. Toupadakis Skouritakis, Director of MediaLab Services at the UC Davis School of Veterinary Medicine, for meticulous figure editing.

## Notes

### Competing Interest Statement

The authors have declared no competing interest.

