## Supplementary material for "A Translational Reference for Green Autofluorescence Imaging in the Rhesus Macaque Eye using the OcuMet Beacon"

### Supplementary Tables

#### Supplementary Table S1. Image quality by region and modality.

| **Quality** | **Sessions** | **Animals** | **Mean** | **SD** |
| --- | --- | --- | --- | --- |
| **Macula (IR)** | 187 | 79 | 83.37 | 10.12 |
| **Macula (GAF)** | 187 | 79 | 66.37 | 6.49 |
| **ONH (IR)** | 208 | 82 | 84.80 | 10.54 |
| **ONH (GAF)** | 209 | 82 | 60.66 | 7.63 |
| **PFB (IR)** | 173 | 78 | 83.00 | 13.53 |
| **PFB (GAF)** | 173 | 78 | 62.95 | 9.39 |

Note: IR, infrared image; GAF, green autofluorescence. “Sessions” counts imaging sessions with analyzable data; “Animals” counts unique animals contributing data.

#### Supplementary Table S2. Green autofluorescence (GAF) intensity by anesthesia method.

| **Region** | **Mean under intramuscular sedation (IM)** | **Mean under general anesthesia (GA)** | ***P*-value** |
| --- | --- | --- | --- |
| **ONH** | 6.85 | 6.34 | 0.4985 |
| **Macula** | 22.04 | 20.39 | 0.9752 |
| **PFB** | 22.42 | 20.84 | 0.6005 |

Note: Values are means; two-sided *P* values show no significant differences between IM and GA in any region. ONH is in dB; macula and PFB are in grey scale units (gsu).

#### Supplementary Table S3. Normative optic nerve head (ONH) green autofluorescence (GAF) intensity by sector and age group in rhesus macaques.

| **Infant & Juvenile (0–5 years, n=21)** | | | | |
| --- | --- | --- | --- | --- |
|  | **Mean** | **SD** | **Low 95 CI** | **Higher CI** |
| **Temporal** | 6.97 | 0.91 | 6.68 | 7.27 |
| **Nasal** | 6.18 | 0.91 | 5.89 | 6.48 |
| **Superior** | 4.63 | 1.11 | 4.29 | 4.98 |
| **Inferior** | 5.14 | 1.21 | 4.75 | 5.53 |
| **Temporal-Superior** | 4.43 | 1.78 | 3.86 | 5.01 |
| **Temporal-Inferior** | 5.48 | 1.34 | 5.05 | 5.92 |
| **Nasal-Superior** | 4.83 | 1.30 | 4.44 | 5.22 |
| **Nasal-Inferior** | 4.80 | 1.64 | 4.28 | 5.33 |
| **Global** | 5.72 | 0.77 | 5.46 | 5.97 |
| **Adult (>5 to <19 years, n=32)** | | | | |
|  | **Mean** | **SD** | **Low 95 CI** | **Higher CI** |
| **Temporal** | 8.39 | 1.53 | 8.05 | 8.73 |
| **Nasal** | 7.04 | 1.57 | 6.70 | 7.38 |
| **Superior** | 5.26 | 1.56 | 4.92 | 5.61 |
| **Inferior** | 5.35 | 1.42 | 5.04 | 5.67 |
| **Temporal-Superior** | 5.54 | 2.06 | 5.09 | 5.99 |
| **Temporal-Inferior** | 5.81 | 1.97 | 5.41 | 6.21 |
| **Nasal-Superior** | 4.99 | 1.80 | 4.60 | 5.39 |
| **Nasal-Inferior** | 4.86 | 2.00 | 4.46 | 5.26 |
| **Global** | 6.51 | 1.31 | 6.22 | 6.80 |
| **Geriatric (≥19 years, n=29)** | | | | |
|  | **Mean** | **SD** | **Low 95 CI** | **Higher CI** |
| **Temporal** | 9.10 | 1.65 | 8.74 | 9.46 |
| **Nasal** | 7.62 | 1.86 | 7.23 | 8.02 |
| **Superior** | 5.67 | 2.02 | 5.24 | 6.11 |
| **Inferior** | 5.60 | 1.81 | 5.21 | 5.99 |
| **Temporal-Superior** | 5.95 | 2.56 | 5.42 | 6.47 |
| **Temporal-Inferior** | 6.06 | 2.43 | 5.64 | 6.47 |
| **Nasal-Superior** | 5.40 | 2.13 | 4.94 | 5.85 |
| **Nasal-Inferior** | 5.14 | 2.56 | 4.69 | 5.59 |
| **Global** | 6.99 | 1.71 | 6.62 | 7.35 |

Note: Values are the mean, standard deviation (SD), and 95% confidence interval (CI); green autofluorescence (GAF) intensity is expressed in decibels (dB); “Global” GAF at the ONH was calculated by averaging the subsectors.

#### Supplementary Table S4. Normative macular green autofluorescence (GAF) intensity and heterogeneity by age group in rhesus macaques.

| **Juvenile-Adult (0–15 years, n=26)** | | | | |
| --- | --- | --- | --- | --- |
|  | **Mean** | **SD** | **Low 95 CI** | **Higher CI** |
| **Intensity** | 17.13 | 6.63 | 15.79 | 18.48 |
| **Heterogeneity** | 0.77 | 0.20 | 0.73 | 0.82 |
| **Mid-Adult (>15 to <19 years, n=25)** | | | | |
|  | **Mean** | **SD** | **Low 95 CI** | **Higher CI** |
| **Intensity** | 22.79 | 5.26 | 20.81 | 24.77 |
| **Heterogeneity** | 0.83 | 0.14 | 0.79 | 0.88 |
| **Geriatric (≥19 years, n=29)** | | | | |
|  | **Mean** | **SD** | **Low 95 CI** | **Higher CI** |
| **Intensity** | 23.84 | 4.07 | 22.80 | 24.89 |
| **Heterogeneity** | 0.84 | 0.21 | 0.79 | 0.90 |

Note: Values are the mean, standard deviation (SD), and 95% confidence interval (CI) of the mean; intensity is reported in grey scale units (gsu), and heterogeneity is a dimensionless index of spatial variability in macular GAF (higher values indicate greater heterogeneity).

#### Supplementary Table S5. Normative peripapillary fundus (PFB) green autofluorescence (GAF) intensity by sector and age group in rhesus macaques.

| **Juvenile-Adult (0–15 years, n=25)** | | | | |
| --- | --- | --- | --- | --- |
|  | **Mean** | **SD** | **Low 95 CI** | **Higher CI** |
| **Horizontal** | 19.83 | 21.08 | 15.41 | 24.25 |
| **Arcuate** | 19.53 | 23.34 | 14.63 | 24.43 |
| **Horizontal-Superior** | 17.74 | 6.30 | 16.36 | 19.12 |
| **Horizontal-Inferior** | 19.42 | 18.26 | 15.58 | 23.25 |
| **Arcuate-Superior** | 16.96 | 6.19 | 15.60 | 18.32 |
| **Arcuate-Inferior** | 18.71 | 15.00 | 15.54 | 21.87 |
| **Global** | 17.38 | 6.24 | 16.01 | 18.75 |
| **Mid-Adult (>15 to <19 years, n=26)** | | | | |
|  | **Mean** | **SD** | **Low 95 CI** | **Higher CI** |
| **Horizontal** | 23.79 | 4.11 | 22.19 | 25.38 |
| **Arcuate** | 22.91 | 4.11 | 21.27 | 24.56 |
| **Horizontal-Superior** | 23.55 | 4.13 | 22.14 | 24.97 |
| **Horizontal-Inferior** | 24.29 | 4.15 | 22.53 | 26.04 |
| **Arcuate-Superior** | 22.52 | 4.02 | 21.13 | 23.90 |
| **Arcuate-Inferior** | 23.50 | 4.36 | 21.64 | 25.37 |
| **Global** | 23.35 | 4.09 | 21.74 | 24.96 |
| **Geriatric (≥19 years, n=28)** | | | | |
|  | **Mean** | **SD** | **Low 95 CI** | **Higher CI** |
| **Horizontal** | 26.50 | 5.74 | 24.69 | 28.31 |
| **Arcuate** | 19.72 | 6.04 | 17.72 | 21.73 |
| **Horizontal-Superior** | 26.63 | 5.76 | 24.62 | 28.64 |
| **Horizontal-Inferior** | 26.42 | 5.75 | 24.74 | 28.09 |
| **Arcuate-Superior** | 20.36 | 6.11 | 18.55 | 22.18 |
| **Arcuate-Inferior** | 24.41 | 6.13 | 21.58 | 27.23 |
| **Global** | 23.11 | 5.89 | 21.21 | 25.02 |

Note: Values are the mean, standard deviation (SD), and 95% confidence interval (CI) of the mean; GAF intensity is reported in grey scale units (gsu). “Global” was calculated by averaging the Horizontal and Arcuate subsectors.

#### Supplementary Table S6. Predictive Factors Associated with ONH GAF Across Subregions.

| **Parameter** | **Predictor** | **Estimate** | **Standard error** | **Low 95 CI** | **High 95 CI** | **P value** | **n** |
| --- | --- | --- | --- | --- | --- | --- | --- |
| **Temp_Inf** | **TRT_CP_S** | -0.0198 | 0.0033 | -0.0261 | -0.0134 | 1.32E-09 | 111 |
| **Nasal** | **TRT_CP_I** | -0.0113 | 0.0021 | -0.0154 | -0.0072 | 6.57E-08 | 111 |
| **Temp_Inf** | **RNFL_CP_S** | -0.0199 | 0.0038 | -0.0273 | -0.0125 | 1.40E-07 | 111 |
| **Global** | **TRT_CP_I** | -0.0093 | 0.0018 | -0.0128 | -0.0058 | 1.82E-07 | 111 |
| **Temp_Inf** | **ONL_CP_I** | -0.0743 | 0.0163 | -0.1063 | -0.0423 | 5.27E-06 | 111 |
| **Temporal** | **TRT_CP_I** | -0.0075 | 0.0018 | -0.0110 | -0.0039 | 3.42E-05 | 111 |
| **Nasal_Sup** | **TRT_CP_I** | -0.0098 | 0.0025 | -0.0147 | -0.0049 | 0.0001 | 111 |
| **Nasal_Sup** | **AWaveTime** | -1.6796 | 0.4290 | -2.5205 | -0.8387 | 0.0001 | 11 |
| **Temp_Inf** | **AWaveTime** | -1.4742 | 0.3967 | -2.2517 | -0.6967 | 0.0002 | 11 |
| **Global** | **RPE_CP_N** | 0.04345 | 0.0124 | 0.0191 | 0.0678 | 0.0005 | 111 |
| **Global** | **OC_CP_I** | -0.0186 | 0.0054 | -0.0292 | -0.0080 | 0.0006 | 111 |
| **Nasal_Inf** | **RNFL_CP_I** | -0.0206 | 0.0060 | -0.0324 | -0.0088 | 0.0006 | 111 |
| **Nasal_Inf** | **RNFL_CP_S** | 0.01423 | 0.0042 | 0.0060 | 0.0225 | 0.0007 | 111 |
| **Temporal** | **RPE_CP_N** | 0.04586 | 0.0136 | 0.0192 | 0.0726 | 0.0008 | 111 |
| **Temporal** | **OC_CP_I** | -0.0187 | 0.0057 | -0.0299 | -0.0075 | 0.0010 | 111 |
| **Nasal_Sup** | **OC_CP_I** | -0.0233 | 0.0072 | -0.0374 | -0.0091 | 0.0013 | 111 |
| **Nasal_Sup** | **CC_mac_NT** | -0.0833 | 0.0261 | -0.1344 | -0.0322 | 0.0014 | 107 |
| **Temporal** | **OS_mac_NT** | -0.1182 | 0.0372 | -0.1911 | -0.0453 | 0.0015 | 107 |
| **Nasal_Sup** | **RPE_CP_N** | 0.05438 | 0.0174 | 0.0203 | 0.0885 | 0.0018 | 111 |
| **Nasal_Inf** | **AWaveTime** | -2.6578 | 0.8557 | -4.3350 | -0.9806 | 0.0019 | 11 |
| **Temp_Inf** | **IPL_CP_T** | -0.0556 | 0.0182 | -0.0913 | -0.0199 | 0.0023 | 111 |
| **Temp_Sup** | **CC_mac_N** | -0.1126 | 0.0373 | -0.1857 | -0.0396 | 0.0025 | 108 |
| **Nasal_Sup** | **TRT_CP_T** | -0.0123 | 0.0041 | -0.0204 | -0.0043 | 0.0026 | 111 |
| **Nasal** | **OC_CP_I** | -0.0196 | 0.0065 | -0.0323 | -0.0068 | 0.0027 | 111 |
| **Nasal** | **OC_CP_S** | -0.0184 | 0.0061 | -0.0304 | -0.0064 | 0.0027 | 111 |
| **Nasal_Sup** | **CC_mac_T** | -0.0649 | 0.0218 | -0.1075 | -0.0222 | 0.0029 | 108 |
| **Global** | **CC_mac_N** | -0.0739 | 0.0251 | -0.1231 | -0.0247 | 0.0032 | 108 |
| **Temporal** | **ONL_CP_I** | -0.0323 | 0.0111 | -0.0541 | -0.0105 | 0.0037 | 111 |
| **Temp_Inf** | **RPE_fov** | 0.03583 | 0.0124 | 0.0114 | 0.0602 | 0.0040 | 108 |
| **Global** | **OC_CP_S** | -0.0147 | 0.0052 | -0.0249 | -0.0046 | 0.0042 | 111 |
| **Nasal_Inf** | **IS_mac_N** | -0.0522 | 0.0183 | -0.0881 | -0.0164 | 0.0043 | 108 |
| **Nasal** | **CC_mac_N** | -0.0865 | 0.0303 | -0.1460 | -0.0271 | 0.0043 | 108 |
| **Nasal_Inf** | **TRT_CP_I** | -0.0121 | 0.0043 | -0.0205 | -0.0037 | 0.0046 | 111 |
| **Nasal_Inf** | **CC_mac_N** | -0.103 | 0.0366 | -0.1746 | -0.0313 | 0.0049 | 108 |
| **Temporal** | **OS_CP_N** | -0.0529 | 0.0188 | -0.0897 | -0.0161 | 0.0049 | 111 |
| **Temporal** | **IPL_CP_T** | -0.0385 | 0.0138 | -0.0656 | -0.0115 | 0.0053 | 111 |
| **Nasal_Inf** | **ONL_CP_S** | -0.0564 | 0.0203 | -0.0962 | -0.0166 | 0.0055 | 111 |
| **Temp_Sup** | **OPL_CP_T** | -0.0936 | 0.0338 | -0.1598 | -0.0273 | 0.0056 | 111 |
| **Nasal_Inf** | **TRT_CP_S** | 0.01239 | 0.0045 | 0.0036 | 0.0212 | 0.0059 | 111 |
| **Temp_Inf** | **IPL_CP_S** | -0.0546 | 0.0199 | -0.0935 | -0.0156 | 0.0060 | 111 |
| **Temp_Sup** | **ONL_CP_S** | -0.048 | 0.0176 | -0.0825 | -0.0136 | 0.0063 | 111 |
| **Nasal_Inf** | **OS_mac_N** | -0.0969 | 0.0357 | -0.1669 | -0.0270 | 0.0066 | 108 |
| **Temp_Inf** | **IS_CP_S** | -0.1549 | 0.0572 | -0.2670 | -0.0427 | 0.0068 | 111 |
| **Nasal** | **GCL_mac_NT** | -0.0698 | 0.0259 | -0.1206 | -0.0190 | 0.0071 | 107 |
| **Nasal** | **TRT_mac_NT** | -0.035 | 0.0130 | -0.0606 | -0.0095 | 0.0071 | 107 |
| **Global** | **CC_mac_NT** | -0.0621 | 0.0231 | -0.1074 | -0.0169 | 0.0072 | 107 |
| **Temporal** | **TRT_mac_NT** | -0.0253 | 0.0095 | -0.0438 | -0.0067 | 0.0076 | 107 |
| **Nasal** | **OS_mac_NT** | -0.1051 | 0.0398 | -0.1831 | -0.0270 | 0.0083 | 107 |
| **Temporal** | **CC_mac_N** | -0.0689 | 0.0264 | -0.1207 | -0.0172 | 0.0090 | 108 |
| **Nasal_Inf** | **OPL_CP_T** | -0.0996 | 0.0385 | -0.1751 | -0.0241 | 0.0097 | 111 |
| **Temporal** | **ONL_CP_S** | -0.0288 | 0.0111 | -0.0506 | -0.0069 | 0.0099 | 111 |
| **Temp_Inf** | **ONL_CP_T** | -0.0498 | 0.0193 | -0.0877 | -0.0119 | 0.0100 | 111 |
| **Temp_Sup** | **OS_CP_N** | -0.0737 | 0.0287 | -0.1300 | -0.0174 | 0.0103 | 111 |
| **Temp_Sup** | **TRT_CP_I** | -0.0109 | 0.0043 | -0.0193 | -0.0026 | 0.0104 | 111 |
| **Nasal_Inf** | **OC_CP_S** | -0.0188 | 0.0073 | -0.0332 | -0.0044 | 0.0104 | 111 |
| **Nasal_Sup** | **TRT_fov** | 0.00592 | 0.0023 | 0.0014 | 0.0105 | 0.0107 | 108 |
| **Temporal** | **OC_CP_N** | -0.0151 | 0.0060 | -0.0269 | -0.0034 | 0.0113 | 111 |
| **Nasal_Sup** | **ONL_CP_I** | -0.0386 | 0.0154 | -0.0688 | -0.0084 | 0.0122 | 111 |
| **Temporal** | **GCL_mac_NT** | -0.0555 | 0.0222 | -0.0990 | -0.0120 | 0.0124 | 107 |
| **Temporal** | **OC_CP_S** | -0.0144 | 0.0058 | -0.0258 | -0.0031 | 0.0126 | 111 |
| **Global** | **OC_CP_N** | -0.0134 | 0.0054 | -0.0240 | -0.0028 | 0.0132 | 111 |
| **Temporal** | **ONL_CP_T** | -0.0396 | 0.0161 | -0.0712 | -0.0080 | 0.0139 | 111 |
| **Nasal_Inf** | **RPE_fov** | -0.0394 | 0.0161 | -0.0709 | -0.0079 | 0.0141 | 108 |
| **Global** | **OS_mac_NT** | -0.0935 | 0.0381 | -0.1682 | -0.0187 | 0.0142 | 107 |
| **Global** | **GCL_mac_NT** | -0.0594 | 0.0243 | -0.1069 | -0.0118 | 0.0144 | 107 |
| **Nasal_Inf** | **OC_CP_I** | -0.0205 | 0.0084 | -0.0369 | -0.0040 | 0.0146 | 111 |
| **Global** | **TRT_mac_NT** | -0.0268 | 0.0110 | -0.0484 | -0.0051 | 0.0153 | 107 |
| **Temporal** | **IS_CP_S** | -0.0885 | 0.0366 | -0.1602 | -0.0168 | 0.0155 | 111 |
| **Temp_Sup** | **OC_CP_S** | -0.0185 | 0.0077 | -0.0335 | -0.0035 | 0.0156 | 111 |
| **Global** | **CC_CP_T** | -0.0811 | 0.0338 | -0.1473 | -0.0149 | 0.0163 | 111 |
| **Global** | **IPL_CP_T** | -0.0332 | 0.0139 | -0.0603 | -0.0060 | 0.0166 | 111 |
| **Global** | **AWaveTime** | -0.9087 | 0.3819 | -1.6572 | -0.1603 | 0.0173 | 11 |
| **Global** | **ONL_CP_T** | -0.0387 | 0.0164 | -0.0708 | -0.0065 | 0.0184 | 111 |
| **Temporal** | **CC_mac_NT** | -0.0578 | 0.0246 | -0.1061 | -0.0096 | 0.0188 | 107 |
| **Temp_Inf** | **GCL_mac_NT** | -0.0763 | 0.0326 | -0.1403 | -0.0124 | 0.0193 | 107 |
| **Nasal** | **ONL_CP_T** | -0.0441 | 0.0189 | -0.0811 | -0.0071 | 0.0196 | 111 |
| **Nasal** | **GCL_mac_T** | -0.0479 | 0.0205 | -0.0881 | -0.0076 | 0.0198 | 108 |
| **Global** | **IS_CP_S** | -0.1001 | 0.0430 | -0.1845 | -0.0158 | 0.0200 | 111 |
| **Global** | **OS_CP_N** | -0.0444 | 0.0192 | -0.0820 | -0.0068 | 0.0205 | 111 |
| **Temp_Sup** | **CC_mac_NT** | -0.0741 | 0.0320 | -0.1369 | -0.0113 | 0.0207 | 107 |
| **Temp_Sup** | **OS_mac_N** | -0.0892 | 0.0386 | -0.1649 | -0.0136 | 0.0208 | 108 |
| **Temporal** | **IS_mac_NT** | -0.0952 | 0.0413 | -0.1761 | -0.0144 | 0.0210 | 107 |
| **Nasal** | **CC_mac_NT** | -0.0646 | 0.0280 | -0.1196 | -0.0097 | 0.0211 | 107 |
| **Temp_Sup** | **IS_CP_S** | -0.1239 | 0.0537 | -0.2292 | -0.0186 | 0.0211 | 111 |
| **Temp_Sup** | **OC_mac_N** | -0.0083 | 0.0036 | -0.0154 | -0.0012 | 0.0212 | 108 |
| **Nasal** | **CC_CP_T** | -0.1045 | 0.0457 | -0.1940 | -0.0151 | 0.0220 | 111 |
| **Global** | **CC_mac_T** | -0.0362 | 0.0158 | -0.0671 | -0.0052 | 0.0220 | 108 |
| **Temp_Sup** | **IS_mac_N** | -0.047 | 0.0206 | -0.0875 | -0.0066 | 0.0226 | 108 |
| **Temp_Sup** | **OPL_CP_I** | -0.101 | 0.0447 | -0.1885 | -0.0135 | 0.0237 | 111 |
| **Temp_Sup** | **OC_mac_T** | -0.0081 | 0.0036 | -0.0152 | -0.0010 | 0.0252 | 108 |
| **Temporal** | **CC_CP_T** | -0.0597 | 0.0267 | -0.1121 | -0.0073 | 0.0257 | 111 |
| **Nasal_Inf** | **RPE_CP_N** | 0.0558 | 0.0250 | 0.0068 | 0.1048 | 0.0257 | 111 |
| **Global** | **OPL_CP_T** | -0.0537 | 0.0243 | -0.1013 | -0.0062 | 0.0267 | 111 |
| **Nasal_Sup** | **GCL_mac_NT** | -0.0728 | 0.0330 | -0.1375 | -0.0081 | 0.0275 | 107 |
| **Nasal_Sup** | **ONL_CP_T** | -0.0445 | 0.0203 | -0.0842 | -0.0048 | 0.0280 | 111 |
| **Temporal** | **OPL_CP_I** | -0.0608 | 0.0277 | -0.1151 | -0.0064 | 0.0284 | 111 |
| **Nasal_Inf** | **OC_CP_N** | -0.0164 | 0.0075 | -0.0310 | -0.0017 | 0.0284 | 111 |
| **Temp_Sup** | **OC_CP_I** | -0.018 | 0.0082 | -0.0341 | -0.0019 | 0.0287 | 111 |
| **Temp_Sup** | **RPE_CP_N** | 0.05214 | 0.0239 | 0.0053 | 0.0990 | 0.0291 | 111 |
| **Nasal_Sup** | **ONL_fov** | 0.01191 | 0.0056 | 0.0010 | 0.0228 | 0.0326 | 108 |
| **Nasal_Inf** | **TRT_CP_T** | -0.0081 | 0.0038 | -0.0156 | -0.0007 | 0.0330 | 111 |
| **Temporal** | **OS_mac_T** | -0.0581 | 0.0273 | -0.1115 | -0.0046 | 0.0332 | 108 |
| **Nasal_Sup** | **TRT_CP_S** | -0.0085 | 0.0040 | -0.0163 | -0.0007 | 0.0336 | 111 |
| **Nasal_Sup** | **RNFL_CP_S** | -0.0092 | 0.0044 | -0.0178 | -0.0007 | 0.0344 | 111 |
| **Nasal** | **IS_CP_S** | -0.1048 | 0.0499 | -0.2026 | -0.0070 | 0.0357 | 111 |
| **Nasal** | **IPL_CP_T** | -0.0314 | 0.0150 | -0.0607 | -0.0021 | 0.0359 | 111 |
| **Temp_Sup** | **OS_CP_T** | -0.0563 | 0.0271 | -0.1094 | -0.0031 | 0.0379 | 111 |
| **Temporal** | **INL_CP_N** | 0.04159 | 0.0201 | 0.0022 | 0.0810 | 0.0386 | 111 |
| **Temp_Inf** | **IS_fov** | 0.02922 | 0.0142 | 0.0014 | 0.0571 | 0.0398 | 108 |
| **Nasal** | **OC_CP_N** | -0.013 | 0.0063 | -0.0254 | -0.0006 | 0.0401 | 111 |
| **Temp_Inf** | **TRT_fov** | 0.00464 | 0.0023 | 0.0002 | 0.0091 | 0.0402 | 108 |
| **Nasal_Sup** | **CC_mac_N** | -0.063 | 0.0307 | -0.1232 | -0.0027 | 0.0405 | 108 |
| **Temporal** | **CC_mac_T** | -0.0336 | 0.0165 | -0.0659 | -0.0013 | 0.0414 | 108 |
| **Temp_Sup** | **OC_mac_NT** | -0.008 | 0.0039 | -0.0156 | -0.0003 | 0.0418 | 107 |
| **Nasal_Inf** | **OS_CP_S** | -0.0649 | 0.0319 | -0.1274 | -0.0024 | 0.0419 | 111 |
| **Temp_Inf** | **CC_CP_N** | 0.0335 | 0.0165 | 0.0011 | 0.0659 | 0.0425 | 111 |
| **Nasal** | **OPL_CP_I** | -0.0726 | 0.0359 | -0.1430 | -0.0023 | 0.0429 | 111 |
| **Temp_Inf** | **ONL_fov** | 0.01085 | 0.0054 | 0.0003 | 0.0214 | 0.0439 | 108 |
| **Nasal** | **TRT_CP_T** | -0.0118 | 0.0059 | -0.0233 | -0.0003 | 0.0442 | 111 |
| **Nasal_Sup** | **CC_CP_T** | -0.1092 | 0.0544 | -0.2159 | -0.0026 | 0.0448 | 111 |
| **Nasal_Sup** | **OC_CP_S** | -0.0135 | 0.0067 | -0.0267 | -0.0003 | 0.0453 | 111 |
| **Nasal_Inf** | **OPL_mac_N** | -0.0467 | 0.0233 | -0.0924 | -0.0009 | 0.0456 | 108 |
| **Global** | **TRT_CP_T** | -0.0093 | 0.0047 | -0.0184 | -0.0002 | 0.0458 | 111 |
| **Temporal** | **GCL_fov** | -0.0311 | 0.0156 | -0.0616 | -0.0005 | 0.0463 | 108 |
| **Nasal_Sup** | **OC_mac_NT** | -0.0067 | 0.0034 | -0.0133 | -0.0001 | 0.0479 | 107 |
| **Global** | **OPL_CP_I** | -0.0639 | 0.0324 | -0.1274 | -0.0005 | 0.0482 | 111 |
| **Temp_Sup** | **CC_CP_S** | -0.116 | 0.0588 | -0.2313 | -0.0007 | 0.0486 | 111 |
| **Nasal_Sup** | **TRT_mac_N** | 0.00252 | 0.0013 | 0.0000 | 0.0050 | 0.0492 | 108 |

Notes. Abbreviations follow the convention layer_region_location; Layer codes, TRT, total retinal thickness, RNFL, retinal nerve fiber layer, GCL, ganglion cell layer, IPL, inner plexiform layer, INL, inner nuclear layer, OPL, outer plexiform layer, ONL, outer nuclear layer, IS, photoreceptor inner segments, OS, photoreceptor outer segments, RPE, retinal pigment epithelium, GCC, ganglion cell complex = RNFL + GCL + IPL, OC, outer choroid, IS+OS, inner + outer segment complex; Region codes, CP, circumpapillary ring scan, mac, macular parafovea, fov, foveal center; Location codes, S, superior, I, inferior, N, nasal, T, temporal; For macular measures, _N and _T indicate the single parafoveal sites 1.5 mm nasal or temporal to the foveal center, and _NT indicates the mean of those two sites; Sector labels, Global, mean across the entire 360 degree measurement, Temporal and Nasal, hemifields on the ring or the single temporal or nasal macular site, Temp_Sup, temporal superior octant, Temp_Inf, temporal inferior octant, Nasal_Sup, nasal superior octant, Nasal_Inf, nasal inferior octant; AWaveTime, full field ERG a wave implicit time; Thickness, micrometers; time, milliseconds.

#### Supplementary Table S7. Predictors of Macular GAF Intensity and Heterogeneity

| **Parameter** | **Predictor** | **Estimate** | **Standard error** | **Low 95 CI** | **High 95 CI** | **P value** | **n** |
| --- | --- | --- | --- | --- | --- | --- | --- |
| **Mac_SI** | **IOP** | -0.6599 | 0.1562 | -0.9661 | -0.3537 | 2.40E-05 | 135 |
| **MacRet** | **OS_mac_NT** | -0.533 | 0.1384 | -0.8042 | -0.2618 | 0.0001 | 101 |
| **MacRet** | **OC_fov** | -0.0465 | 0.0137 | -0.0735 | -0.0196 | 0.0007 | 102 |
| **MacRet** | **OS_mac_T** | -0.3707 | 0.1188 | -0.6036 | -0.1378 | 0.0018 | 102 |
| **MacRet** | **RPE_mac_T** | 0.20218 | 0.0679 | 0.0690 | 0.3353 | 0.0029 | 102 |
| **MacRet** | **IOP** | 0.25425 | 0.0876 | 0.0826 | 0.4259 | 0.0037 | 135 |
| **Mac_SI** | **RPE_CP_T** | 0.00573 | 0.0022 | 0.0015 | 0.0100 | 0.0084 | 106 |
| **MacRet** | **OC_mac_T** | -0.0297 | 0.0113 | -0.0519 | -0.0076 | 0.0086 | 102 |
| **MacRet** | **CC_mac_N** | -0.209 | 0.0804 | -0.3665 | -0.0515 | 0.0093 | 102 |
| **MacRet** | **GCL_fov** | -0.1295 | 0.0500 | -0.2275 | -0.0315 | 0.0096 | 102 |
| **MacRet** | **GCL_mac_T** | 0.1455 | 0.0572 | 0.0334 | 0.2576 | 0.0110 | 102 |
| **MacRet** | **OC_mac_N** | -0.0253 | 0.0101 | -0.0452 | -0.0054 | 0.0126 | 102 |
| **Mac_SI** | **RPE_CP_I** | 0.00725 | 0.0030 | 0.0014 | 0.0131 | 0.0159 | 106 |
| **Mac_SI** | **RPE_CP_S** | 0.00884 | 0.0037 | 0.0016 | 0.0161 | 0.0170 | 106 |
| **MacRet** | **OC_mac_NT** | -0.03 | 0.0126 | -0.0547 | -0.0052 | 0.0175 | 101 |
| **Mac_SI** | **RPE_CP_N** | 0.00406 | 0.0017 | 0.0007 | 0.0075 | 0.0194 | 106 |
| **MacRet** | **CC_mac_NT** | -0.2165 | 0.0946 | -0.4019 | -0.0311 | 0.0221 | 101 |
| **Mac_SI** | **GCL_CP_I** | 0.00608 | 0.0028 | 0.0006 | 0.0116 | 0.0300 | 106 |
| **MacRet** | **CC_mac_T** | -0.1705 | 0.0787 | -0.3248 | -0.0162 | 0.0303 | 102 |
| **MacRet** | **RPE_mac_NT** | 0.17321 | 0.0808 | 0.0149 | 0.3315 | 0.0320 | 101 |
| **Mac_SI** | **INL_CP_N** | 0.00882 | 0.0041 | 0.0007 | 0.0169 | 0.0324 | 106 |
| **MacRet** | **CC_fov** | -0.0857 | 0.0413 | -0.1666 | -0.0048 | 0.0380 | 102 |
| **Mac_SI** | **OPL_CP_I** | 0.00827 | 0.0041 | 0.0002 | 0.0163 | 0.0444 | 106 |
| **MacRet** | **INL_CP_I** | 0.13469 | 0.0672 | 0.0031 | 0.2663 | 0.0449 | 106 |
| **MacRet** | **ONL_CP_T** | 0.0973 | 0.0492 | 0.0008 | 0.1938 | 0.0482 | 106 |

Note MacRet, flavoprotein fluorescence (GAF) intensity in grey scale units (gsu); Mac_SI, GAF heterogeneity (unitless index); Predictors, use the convention layer_region_location; Layers, OS, photoreceptor outer segments, IS, inner segments, ONL, outer nuclear layer, INL, inner nuclear layer, IPL, inner plexiform layer, OPL, outer plexiform layer, GCL, ganglion cell layer, RPE, retinal pigment epithelium, GCC, ganglion cell complex = RNFL + GCL + IPL, OC, outer choroid; Regions, mac, macular parafovea, fov, foveal center, CP, circumpapillary ring scan; Locations and sectors, N, nasal, T, temporal, S, superior, I, inferior; for macular measures, _N and _T denote the single parafoveal sites 1.5 mm nasal or temporal to the foveal center, and _NT denotes the mean of those two sites; for CP measures, sector letters denote the corresponding hemifield, and Temp_Sup, Temp_Inf, Nasal_Sup, and Nasal_Inf denote the temporal superior, temporal inferior, nasal superior, and nasal inferior octants; IOP, intraocular pressure (mmHg); Thickness values, micrometers.

#### Supplementary Table S8. Predictors of PFB GAF Intensity

| **Parameter** | **Predictor** | Estimate | Standard error | Low 95 CI | High 95 CI | P value | n |
| --- | --- | --- | --- | --- | --- | --- | --- |
| **Sect_AI** | **OS_mac_NT** | -0.5654 | 0.1047 | -0.7707 | -0.3601 | 0.0001 | 95 |
| **Sect_HI** | **OS_mac_NT** | -0.5256 | 0.1060 | -0.7332 | -0.3179 | 0.0001 | 95 |
| **Global** | **OS_mac_NT** | -0.5042 | 0.1018 | -0.7037 | -0.3047 | 0.0001 | 95 |
| **Sect_HS** | **OS_mac_NT** | -0.4851 | 0.1051 | -0.6911 | -0.2790 | 0.0001 | 95 |
| **Sect_AS** | **OS_mac_NT** | -0.4408 | 0.1011 | -0.6388 | -0.2427 | 0.0001 | 95 |
| **Sect_HI** | **OS_mac_T** | -0.3295 | 0.0918 | -0.5094 | -0.1497 | 0.0003 | 96 |
| **Sect_AI** | **RPE_mac_T** | 0.19787 | 0.0575 | 0.0852 | 0.3105 | 0.0006 | 96 |
| **Sect_HI** | **AWaveTime** | 4.40845 | 1.2840 | 1.8919 | 6.9249 | 0.0006 | 10 |
| **Global** | **AWaveTime** | 2.66453 | 0.7783 | 1.1390 | 4.1900 | 0.0006 | 10 |
| **Sect_AI** | **OS_mac_T** | -0.3444 | 0.1009 | -0.5422 | -0.1466 | 0.0006 | 96 |
| **Sect_HS** | **OS_mac_T** | -0.3063 | 0.0903 | -0.4832 | -0.1293 | 0.0007 | 96 |
| **Global** | **OS_mac_T** | -0.3146 | 0.0933 | -0.4974 | -0.1318 | 0.0007 | 96 |
| **Sect_HI** | **OPL_CP_N** | -0.2305 | 0.0699 | -0.3675 | -0.0934 | 0.0010 | 98 |
| **Sect_HI** | **RPE_mac_T** | 0.18774 | 0.0574 | 0.0752 | 0.3003 | 0.0011 | 96 |
| **Global** | **RPE_mac_T** | 0.18014 | 0.0562 | 0.0699 | 0.2904 | 0.0014 | 96 |
| **Sect_HS** | **RPE_mac_T** | 0.18015 | 0.0573 | 0.0678 | 0.2925 | 0.0017 | 96 |
| **Sect_HI** | **OS_mac_N** | -0.2851 | 0.0923 | -0.4660 | -0.1042 | 0.0020 | 96 |
| **Sect_AI** | **OC_fov** | -0.0422 | 0.0139 | -0.0694 | -0.0150 | 0.0024 | 96 |
| **Global** | **OPL_CP_N** | -0.21 | 0.0695 | -0.3463 | -0.0736 | 0.0025 | 98 |
| **Sect_HI** | **INL_CP_I** | 0.19427 | 0.0647 | 0.0675 | 0.3210 | 0.0027 | 98 |
| **Sect_AS** | **OS_mac_T** | -0.2783 | 0.0935 | -0.4616 | -0.0950 | 0.0029 | 96 |
| **Sect_AI** | **RPE_mac_NT** | 0.20821 | 0.0702 | 0.0706 | 0.3458 | 0.0030 | 95 |
| **Sect_HS** | **OC_fov** | -0.0419 | 0.0142 | -0.0697 | -0.0142 | 0.0031 | 96 |
| **Sect_HI** | **OC_fov** | -0.0413 | 0.0140 | -0.0688 | -0.0137 | 0.0033 | 96 |
| **Sect_HS** | **INL_CP_I** | 0.18833 | 0.0641 | 0.0627 | 0.3140 | 0.0033 | 98 |
| **Global** | **OC_fov** | -0.0406 | 0.0139 | -0.0678 | -0.0135 | 0.0034 | 96 |
| **Sect_HS** | **OPL_CP_N** | -0.2136 | 0.0730 | -0.3568 | -0.0705 | 0.0034 | 98 |
| **Sect_AI** | **OS_mac_N** | -0.291 | 0.1006 | -0.4881 | -0.0938 | 0.0038 | 96 |
| **Global** | **INL_CP_I** | 0.18 | 0.0628 | 0.0570 | 0.3030 | 0.0041 | 98 |
| **Sect_HS** | **OS_mac_N** | -0.2593 | 0.0909 | -0.4374 | -0.0811 | 0.0043 | 96 |
| **Sect_AI** | **OPL_CP_N** | -0.2014 | 0.0711 | -0.3406 | -0.0621 | 0.0046 | 98 |
| **Global** | **OS_mac_N** | -0.261 | 0.0922 | -0.4417 | -0.0803 | 0.0046 | 96 |
| **Sect_AS** | **RPE_mac_T** | 0.15481 | 0.0560 | 0.0450 | 0.2646 | 0.0057 | 96 |
| **Sect_HI** | **RPE_mac_NT** | 0.19534 | 0.0709 | 0.0564 | 0.3343 | 0.0059 | 95 |
| **Sect_AI** | **OC_mac_T** | -0.0307 | 0.0112 | -0.0527 | -0.0087 | 0.0062 | 96 |
| **Global** | **RPE_mac_NT** | 0.18495 | 0.0683 | 0.0511 | 0.3188 | 0.0068 | 95 |
| **Sect_AS** | **OC_mac_T** | -0.0282 | 0.0105 | -0.0488 | -0.0076 | 0.0073 | 96 |
| **Sect_AI** | **INL_CP_I** | 0.1816 | 0.0677 | 0.0489 | 0.3143 | 0.0073 | 98 |
| **Sect_AS** | **OC_fov** | -0.037 | 0.0138 | -0.0642 | -0.0099 | 0.0074 | 96 |
| **Sect_AI** | **CC_mac_N** | -0.2007 | 0.0752 | -0.3482 | -0.0533 | 0.0076 | 96 |
| **Global** | **OC_mac_T** | -0.0297 | 0.0111 | -0.0515 | -0.0078 | 0.0078 | 96 |
| **Sect_HS** | **OC_mac_T** | -0.0307 | 0.0116 | -0.0533 | -0.0080 | 0.0079 | 96 |
| **Sect_AS** | **OPL_CP_N** | -0.1944 | 0.0736 | -0.3386 | -0.0502 | 0.0082 | 98 |
| **Sect_HS** | **RPE_mac_NT** | 0.18314 | 0.0697 | 0.0466 | 0.3197 | 0.0086 | 95 |
| **Sect_HS** | **CC_fov** | -0.0894 | 0.0348 | -0.1576 | -0.0213 | 0.0101 | 96 |
| **Sect_AS** | **INL_CP_I** | 0.15578 | 0.0606 | 0.0370 | 0.2745 | 0.0101 | 98 |
| **Sect_HI** | **CC_fov** | -0.0897 | 0.0351 | -0.1584 | -0.0209 | 0.0106 | 96 |
| **Sect_AI** | **CC_mac_NT** | -0.2132 | 0.0844 | -0.3785 | -0.0478 | 0.0115 | 95 |
| **Sect_AI** | **CC_fov** | -0.0805 | 0.0319 | -0.1429 | -0.0180 | 0.0116 | 96 |
| **Global** | **BWaveTime** | 0.22487 | 0.0891 | 0.0502 | 0.3995 | 0.0116 | 127 |
| **Global** | **CC_fov** | -0.0841 | 0.0334 | -0.1496 | -0.0185 | 0.0119 | 96 |
| **Global** | **CC_mac_N** | -0.1864 | 0.0741 | -0.3316 | -0.0411 | 0.0119 | 96 |
| **Sect_HI** | **OC_mac_T** | -0.029 | 0.0116 | -0.0517 | -0.0064 | 0.0120 | 96 |
| **Sect_AS** | **IOP** | 0.25492 | 0.1020 | 0.0550 | 0.4548 | 0.0124 | 127 |
| **Sect_AI** | **CC_mac_T** | -0.1794 | 0.0727 | -0.3219 | -0.0370 | 0.0136 | 96 |
| **Sect_HI** | **IOP** | 0.21807 | 0.0884 | 0.0449 | 0.3912 | 0.0136 | 127 |
| **Global** | **CC_mac_NT** | -0.1992 | 0.0814 | -0.3587 | -0.0397 | 0.0144 | 95 |
| **Sect_AS** | **CC_mac_NT** | -0.1784 | 0.0731 | -0.3218 | -0.0351 | 0.0147 | 95 |
| **Sect_HS** | **CC_mac_N** | -0.1857 | 0.0768 | -0.3362 | -0.0352 | 0.0156 | 96 |
| **Sect_AS** | **CC_mac_N** | -0.1703 | 0.0705 | -0.3085 | -0.0320 | 0.0158 | 96 |
| **Global** | **CC_mac_T** | -0.1696 | 0.0704 | -0.3075 | -0.0316 | 0.0160 | 96 |
| **Sect_AI** | **OC_mac_N** | -0.0219 | 0.0091 | -0.0397 | -0.0041 | 0.0161 | 96 |
| **Sect_HI** | **CC_mac_N** | -0.1889 | 0.0785 | -0.3428 | -0.0349 | 0.0162 | 96 |
| **Sect_HS** | **CC_mac_NT** | -0.1997 | 0.0832 | -0.3627 | -0.0367 | 0.0164 | 95 |
| **Sect_AS** | **CC_mac_T** | -0.1472 | 0.0614 | -0.2675 | -0.0268 | 0.0165 | 96 |
| **Sect_AI** | **OC_mac_NT** | -0.028 | 0.0117 | -0.0509 | -0.0050 | 0.0169 | 95 |
| **Sect_HS** | **CC_mac_T** | -0.172 | 0.0723 | -0.3138 | -0.0303 | 0.0173 | 96 |
| **Sect_HS** | **AWaveTime** | 2.3395 | 0.9893 | 0.4006 | 4.2784 | 0.0180 | 10 |
| **Sect_HS** | **OC_mac_N** | -0.0219 | 0.0093 | -0.0401 | -0.0036 | 0.0189 | 96 |
| **Sect_HS** | **ONL_CP_T** | 0.11582 | 0.0494 | 0.0191 | 0.2126 | 0.0190 | 98 |
| **Sect_AI** | **IOP** | 0.21126 | 0.0901 | 0.0346 | 0.3879 | 0.0191 | 127 |
| **Sect_HI** | **CC_mac_NT** | -0.2056 | 0.0883 | -0.3787 | -0.0325 | 0.0199 | 95 |
| **Sect_HS** | **IOP** | 0.21194 | 0.0912 | 0.0332 | 0.3907 | 0.0201 | 127 |
| **Sect_AS** | **OS_mac_N** | -0.2086 | 0.0899 | -0.3849 | -0.0324 | 0.0203 | 96 |
| **Sect_AS** | **GCL_mac_T** | 0.12213 | 0.0527 | 0.0188 | 0.2255 | 0.0206 | 96 |
| **Global** | **OC_mac_NT** | -0.0265 | 0.0115 | -0.0491 | -0.0040 | 0.0212 | 95 |
| **Sect_HI** | **CC_mac_T** | -0.1796 | 0.0779 | -0.3322 | -0.0269 | 0.0212 | 96 |
| **Sect_HS** | **OC_mac_NT** | -0.0276 | 0.0120 | -0.0510 | -0.0041 | 0.0214 | 95 |
| **Global** | **OC_mac_N** | -0.0206 | 0.0089 | -0.0381 | -0.0031 | 0.0214 | 96 |
| **Sect_AS** | **GCL_mac_NT** | 0.16182 | 0.0706 | 0.0235 | 0.3001 | 0.0218 | 95 |
| **Sect_AS** | **RPE_mac_NT** | 0.15312 | 0.0668 | 0.0222 | 0.2840 | 0.0219 | 95 |
| **Sect_HI** | **OC_mac_N** | -0.0216 | 0.0094 | -0.0400 | -0.0031 | 0.0220 | 96 |
| **Sect_AS** | **OC_mac_NT** | -0.0244 | 0.0107 | -0.0454 | -0.0035 | 0.0224 | 95 |
| **Sect_AS** | **TRT_mac_T** | 0.01521 | 0.0067 | 0.0021 | 0.0283 | 0.0227 | 96 |
| **Sect_AI** | **TRT_mac_T** | 0.01486 | 0.0065 | 0.0021 | 0.0277 | 0.0229 | 96 |
| **Sect_HI** | **IPL_CP_N** | -0.1332 | 0.0592 | -0.2491 | -0.0172 | 0.0244 | 98 |
| **Sect_HI** | **ONL_CP_T** | 0.10934 | 0.0487 | 0.0139 | 0.2048 | 0.0247 | 98 |
| **Sect_AI** | **IPL_CP_N** | -0.1359 | 0.0607 | -0.2549 | -0.0168 | 0.0253 | 98 |
| **Sect_AS** | **CC_fov** | -0.0766 | 0.0345 | -0.1443 | -0.0089 | 0.0265 | 96 |
| **Sect_HS** | **IPL_CP_N** | -0.1317 | 0.0595 | -0.2483 | -0.0152 | 0.0268 | 98 |
| **Global** | **IPL_CP_N** | -0.127 | 0.0578 | -0.2403 | -0.0137 | 0.0280 | 98 |
| **Global** | **ONL_CP_T** | 0.10246 | 0.0469 | 0.0106 | 0.1943 | 0.0288 | 98 |
| **Sect_HI** | **OC_mac_NT** | -0.0261 | 0.0120 | -0.0496 | -0.0027 | 0.0289 | 95 |
| **Sect_AS** | **ONL_CP_T** | 0.10507 | 0.0484 | 0.0102 | 0.1999 | 0.0299 | 98 |
| **Global** | **TRT_mac_T** | 0.01413 | 0.0065 | 0.0014 | 0.0269 | 0.0299 | 96 |
| **Sect_HS** | **INL_CP_N** | 0.1829 | 0.0846 | 0.0170 | 0.3488 | 0.0307 | 98 |
| **Sect_HI** | **INL_CP_N** | 0.17766 | 0.0836 | 0.0138 | 0.3415 | 0.0336 | 98 |
| **Global** | **GCL_mac_NT** | 0.14332 | 0.0679 | 0.0103 | 0.2764 | 0.0347 | 95 |
| **Sect_HS** | **TRT_mac_T** | 0.01389 | 0.0066 | 0.0009 | 0.0269 | 0.0360 | 96 |
| **Global** | **GCL_mac_T** | 0.1021 | 0.0488 | 0.0065 | 0.1977 | 0.0364 | 96 |
| **Sect_AI** | **INL_CP_N** | 0.17382 | 0.0835 | 0.0102 | 0.3375 | 0.0374 | 98 |
| **Global** | **INL_CP_N** | 0.17234 | 0.0834 | 0.0088 | 0.3358 | 0.0388 | 98 |
| **Sect_HI** | **IS_fov** | -0.0453 | 0.0221 | -0.0886 | -0.0020 | 0.0403 | 96 |
| **Sect_AS** | **AWaveTime** | 1.9868 | 0.9705 | 0.0847 | 3.8889 | 0.0406 | 10 |
| **Sect_HS** | **GCL_mac_NT** | 0.14518 | 0.0710 | 0.0059 | 0.2844 | 0.0410 | 95 |
| **Sect_AI** | **GCL_mac_T** | 0.0968 | 0.0480 | 0.0028 | 0.1908 | 0.0436 | 96 |
| **Sect_AS** | **OC_mac_N** | -0.017 | 0.0084 | -0.0335 | -0.0004 | 0.0444 | 96 |
| **Sect_AI** | **AWaveTime** | 1.92339 | 0.9689 | 0.0243 | 3.8224 | 0.0471 | 10 |
| **Sect_HI** | **GCL_mac_NT** | 0.13323 | 0.0678 | 0.0004 | 0.2661 | 0.0493 | 95 |

Notes: Papillofoveal bundle (PFB) outcomes, Sect_AS, arcuate superior, Sect_AI, arcuate inferior, Sect_HS, horizontal superior, Sect_HI, horizontal inferior, Global, mean across all PFB sectors; Predictors, use the convention layer_region_location; Layers, OS, photoreceptor outer segments, IS, inner segments, ONL, outer nuclear layer, INL, inner nuclear layer, IPL, inner plexiform layer, OPL, outer plexiform layer, GCL, ganglion cell layer, RPE, retinal pigment epithelium, GCC, ganglion cell complex = RNFL + GCL + IPL, OC, outer choroid; Regions, mac, macular parafovea, fov, foveal center, CP, circumpapillary ring scan; Locations and sectors, N, nasal, T, temporal, S, superior, I, inferior; for macular variables, _N and _T denote the single parafoveal sites 1.5 mm nasal or temporal to the foveal center, and _NT denotes the mean of those two sites; for CP variables, letters denote the corresponding hemifield or octant as applicable; IOP, intraocular pressure (mmHg); AWaveTime, full field ERG a wave implicit time (ms); flavoprotein fluorescence intensity, grey scale units (gsu); thickness values, micrometers.

### Supplementary Figures

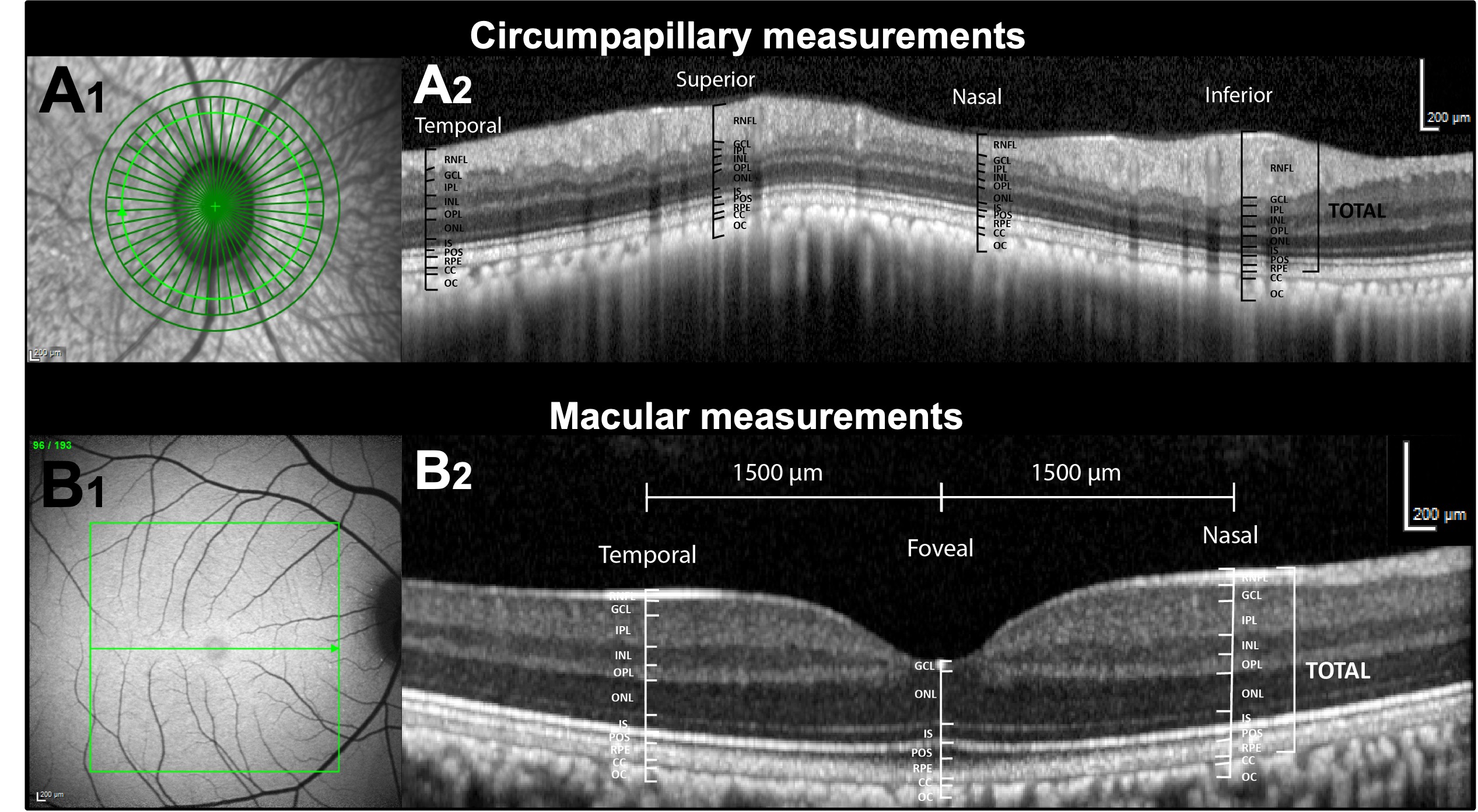

**Supplementary Figure S1. Anatomical locations used for OCT retinal layer measurements.** **A1-A2**, circumpapillary measurements centered on the optic nerve head (ONH). **A1** shows the infrared cSLO with the peripapillary circular scan and segmentation grid; **A2** shows the corresponding circumpapillary B scan with the four sampled locations (temporal, superior, inferior, nasal) and representative layer boundaries used for thickness measurements. **B1-B2**, macular measurements centered on the fovea. **B1** shows the blue autofluorescence cSLO with the macular scan region; **B2** shows the macular B scan with the two parafoveal sampling sites located 1.5 mm temporal and 1.5 mm nasal to the foveal center, together with representative layer boundaries.

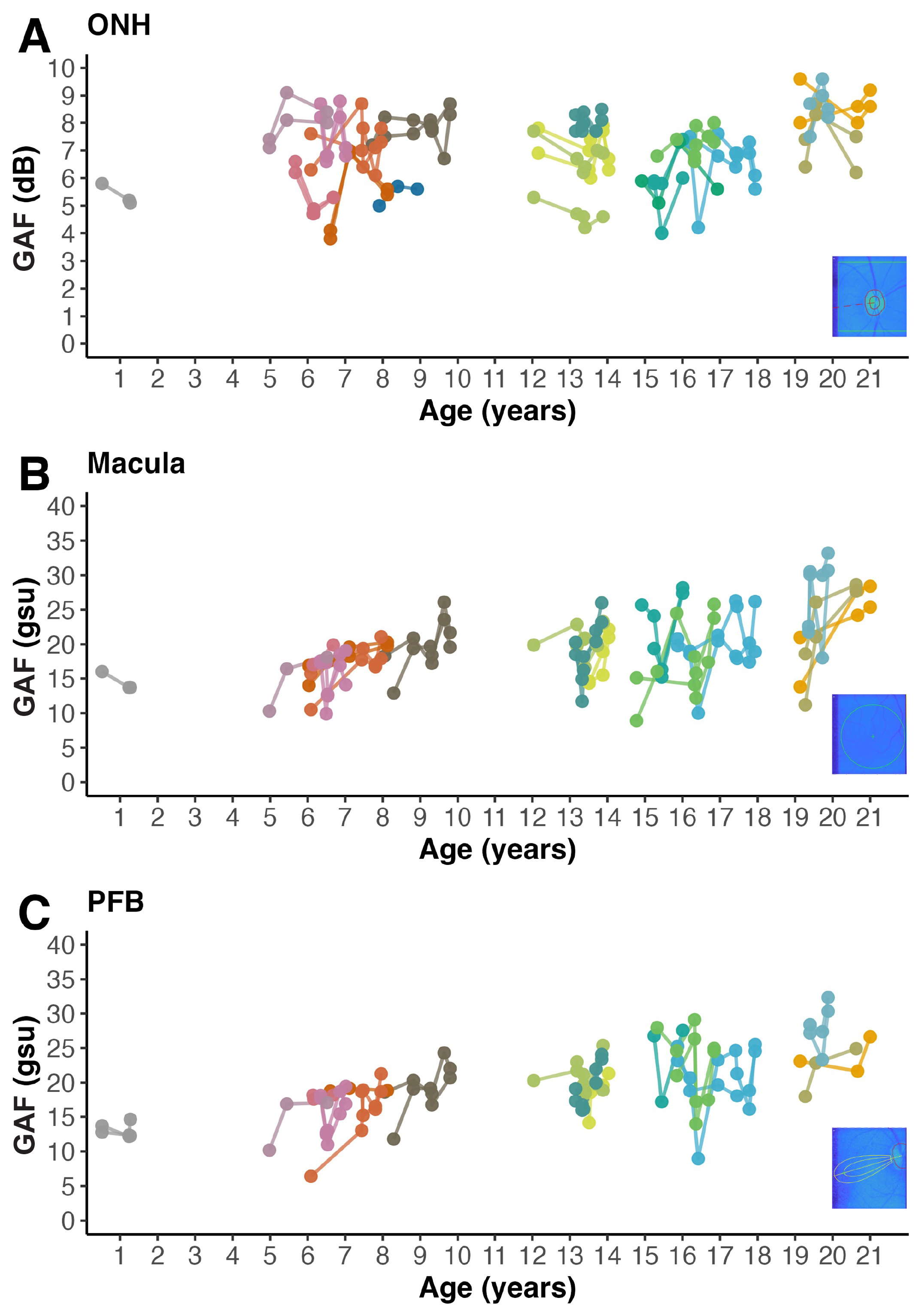

**Supplementary Figure S2. Longitudinal trajectories show low within-eye variability in green autofluorescence (GAF) across sessions in the three retinal regions evaluated**. A, optic nerve head (ONH; dB); B, macula (grey scale units, gsu); C, papillofoveal bundle (PFB; grey scale units, gsu). Each data point represents one eye at a single imaging session. Solid lines connect repeated sessions from the same eye to illustrate the trajectory over time. Only animals with at least three imaging sessions are included, and a distinct color identifies each animal with the same color used across panels. Insets show representative pseudocolor GAF images from the analyzed regions for reference.

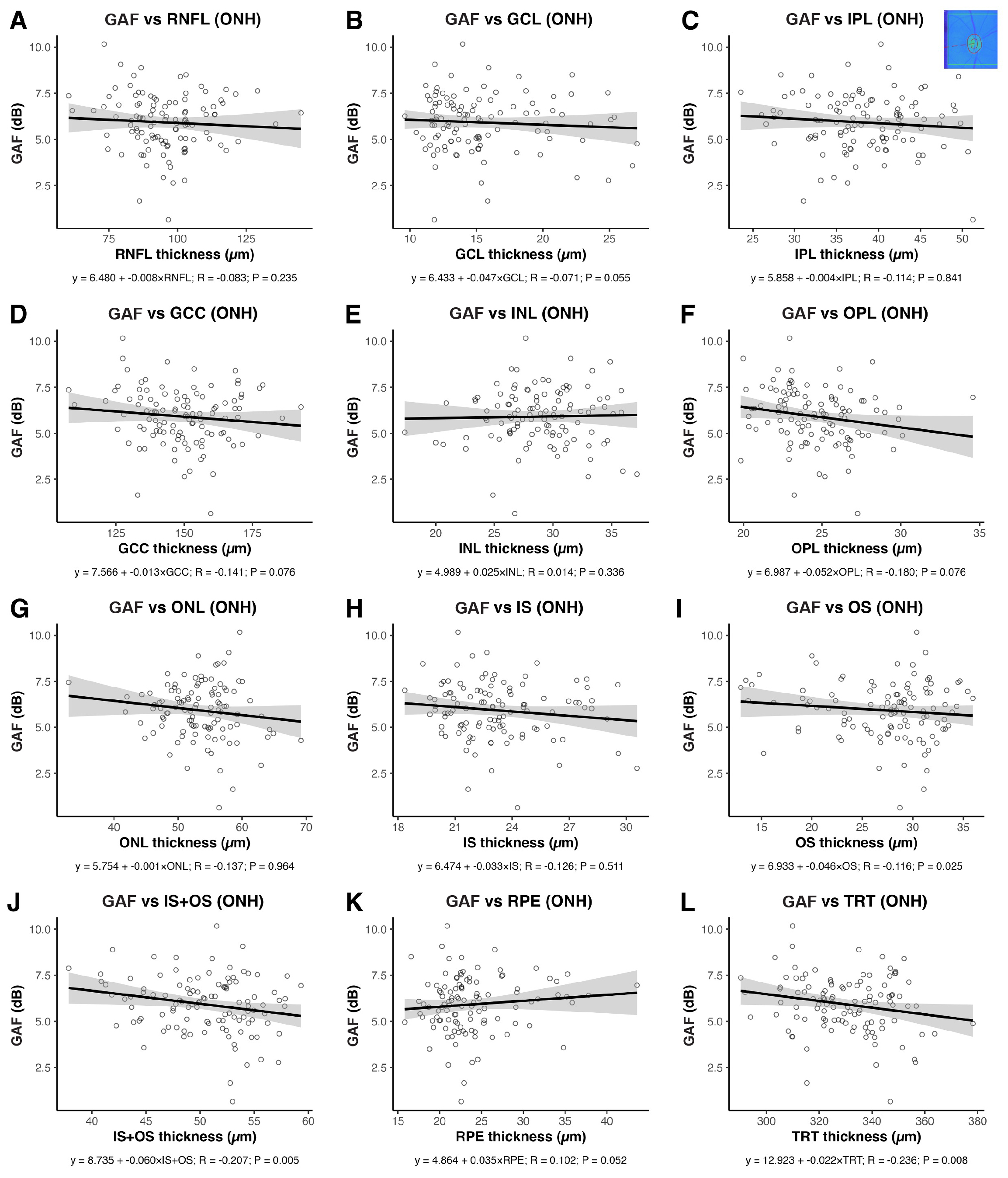

**Supplementary Figure S3. After adjusting for age, optic nerve head (ONH) green autofluorescence (GAF) increased as most neuroretinal layers thinned. A**, retinal nerve fiber layer (RNFL) thickness; **B**, ganglion cell layer (GCL) thickness; **C**, inner plexiform layer (IPL) thickness; **D**, ganglion cell complex (GCC = RNFL + GCL + IPL) thickness; **E**, inner nuclear layer (INL) thickness; **F**, outer plexiform layer (OPL) thickness; **G**, outer nuclear layer (ONL) thickness; **H**, photoreceptor inner segment (IS) thickness; **I,** photoreceptor outer segment (OS) thickness; **J**, IS + OS combination thickness; **K**, retinal pigment epithelium (RPE) thickness; **L**, total retinal thickness (TRT). Retinal layer thickness was measured on the circumpapillary ONH scan as the mean of four locations: superior, inferior, temporal, and nasal. Each data point represents one eye at a single imaging session with OCT and GAF obtained at the same visit. Solid lines show linear fits from mixed models accounting for intereye and intraindividual repeated measures correlation with 95 percent confidence bands. Annotations in each panel report the slope in dB per µm together with the model R and P values. Insets show representative pseudocolor GAF images from the analyzed regions for reference.

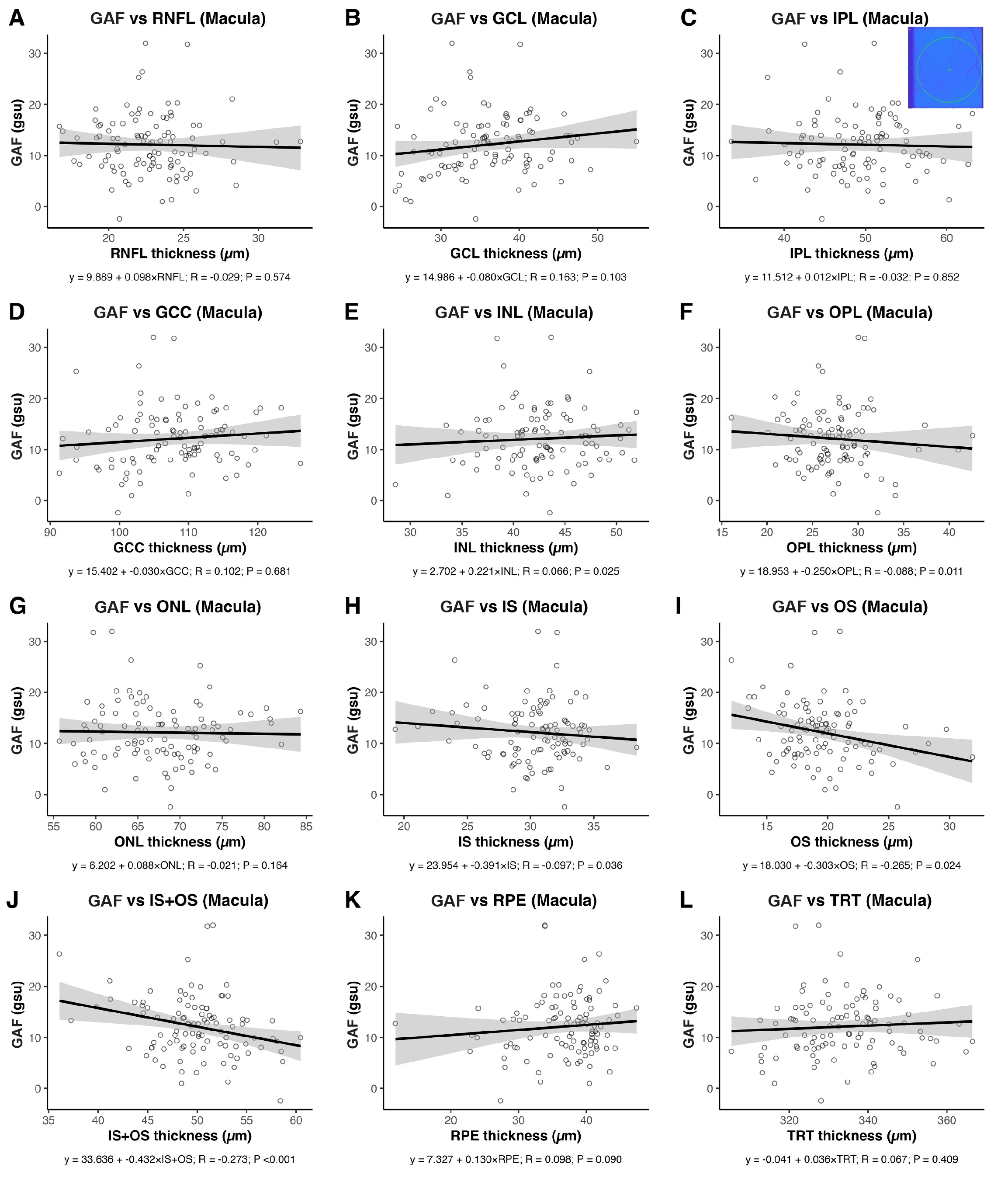

**Supplementary Figure S4. Age-adjusted macular green autofluorescence (GAF) is inversely related to outer retinal thickness and slightly positively related to the inner nuclear layer (INL). A**, retinal nerve fiber layer (RNFL) thickness; **B**, ganglion cell layer (GCL) thickness; **C**, inner plexiform layer (IPL) thickness; **D**, ganglion cell complex (GCC = RNFL + GCL + IPL) thickness; **E**, inner nuclear layer (INL) thickness; **F**, outer plexiform layer (OPL) thickness; **G**, outer nuclear layer (ONL) thickness; **H**, photoreceptor inner segment (IS) thickness; **I,** photoreceptor outer segment (OS) thickness; **J**, IS + OS combination thickness; **K**, retinal pigment epithelium (RPE) thickness; **L**, total retinal thickness (TRT). Retinal layer thickness was measured on the macular OCT scan at two parafoveal sites located 1.5 mm nasal and temporal to the foveal center; the mean of these two measurements was used for analysis. Each point represents one eye at a single visit in which OCT and macular GAF were obtained. Solid lines show linear fits from mixed models accounting for intereye and intraindividual repeated measures correlation within subjects, with 95% confidence bands. Panel annotations report the model slope (in gsu per µm), together with the model R and P values. Insets show representative pseudocolor GAF images from the analyzed regions for reference.

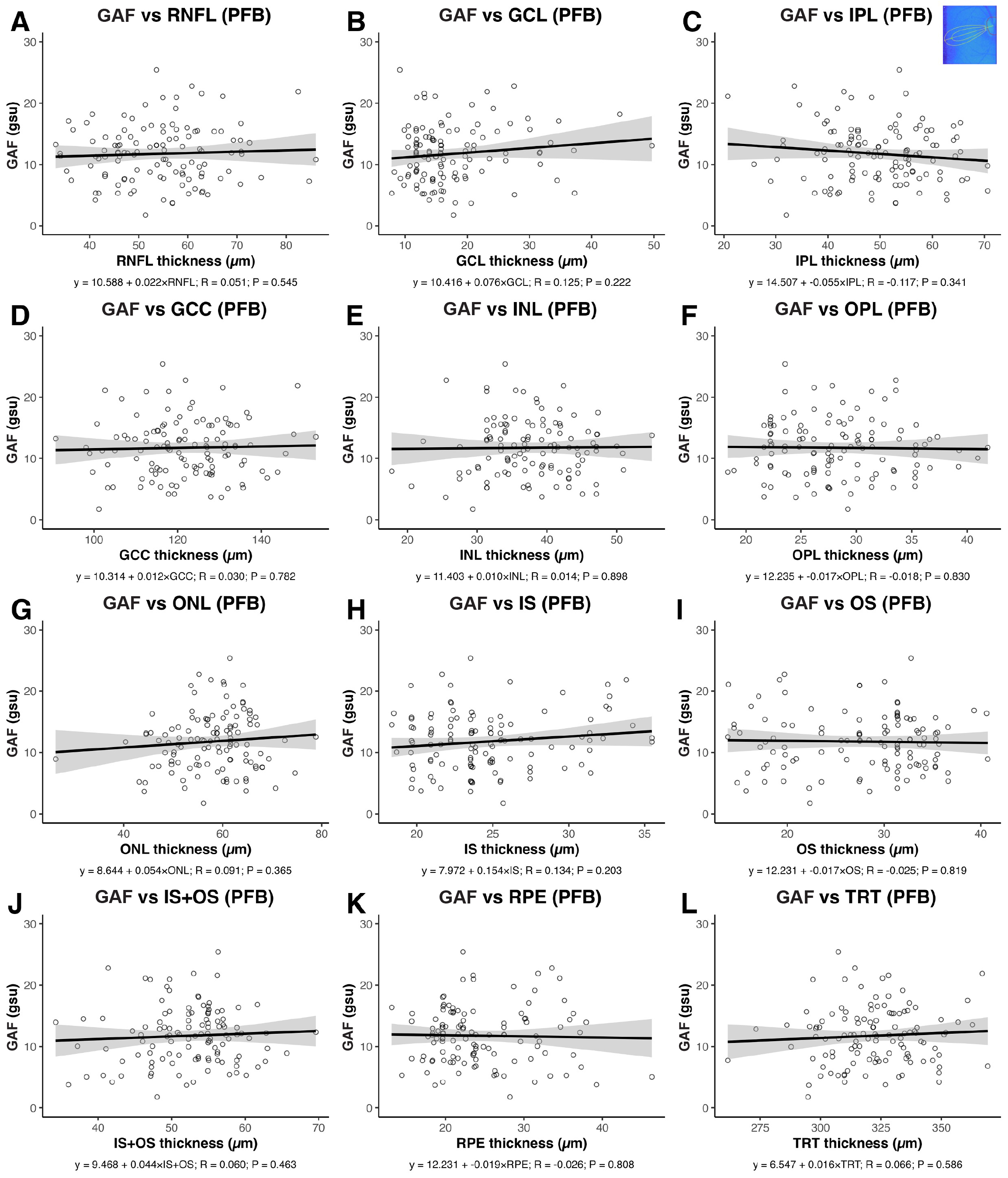

**Supplementary Figure S5. After age adjustment, papillofoveal bundle (PFB) green autofluorescence (GAF) shows only weak, nonsignificant associations with retinal layer thickness. A**, retinal nerve fiber layer (RNFL) thickness; **B**, ganglion cell layer (GCL) thickness; **C**, inner plexiform layer (IPL) thickness; **D**, ganglion cell complex (GCC = RNFL + GCL + IPL) thickness; **E**, inner nuclear layer (INL) thickness; **F**, outer plexiform layer (OPL) thickness; **G**, outer nuclear layer (ONL) thickness; **H**, photoreceptor inner segment (IS) thickness; **I,** photoreceptor outer segment (OS) thickness; **J**, IS + OS combination thickness; **K**, retinal pigment epithelium (RPE) thickness; **L**, total retinal thickness (TRT). Retinal layer thickness was obtained from the circumpapillary ONH scan at a single temporal location. Each data point represents one eye at a single imaging session with OCT and GAF obtained at the same visit. Solid lines show linear fits from mixed models accounting for intereye and intraindividual repeated measures correlation with 95 percent confidence bands. Annotations in each panel report the slope in gsu per µm together with the model R and P values. Insets show representative pseudocolor GAF images from the analyzed regions for reference.

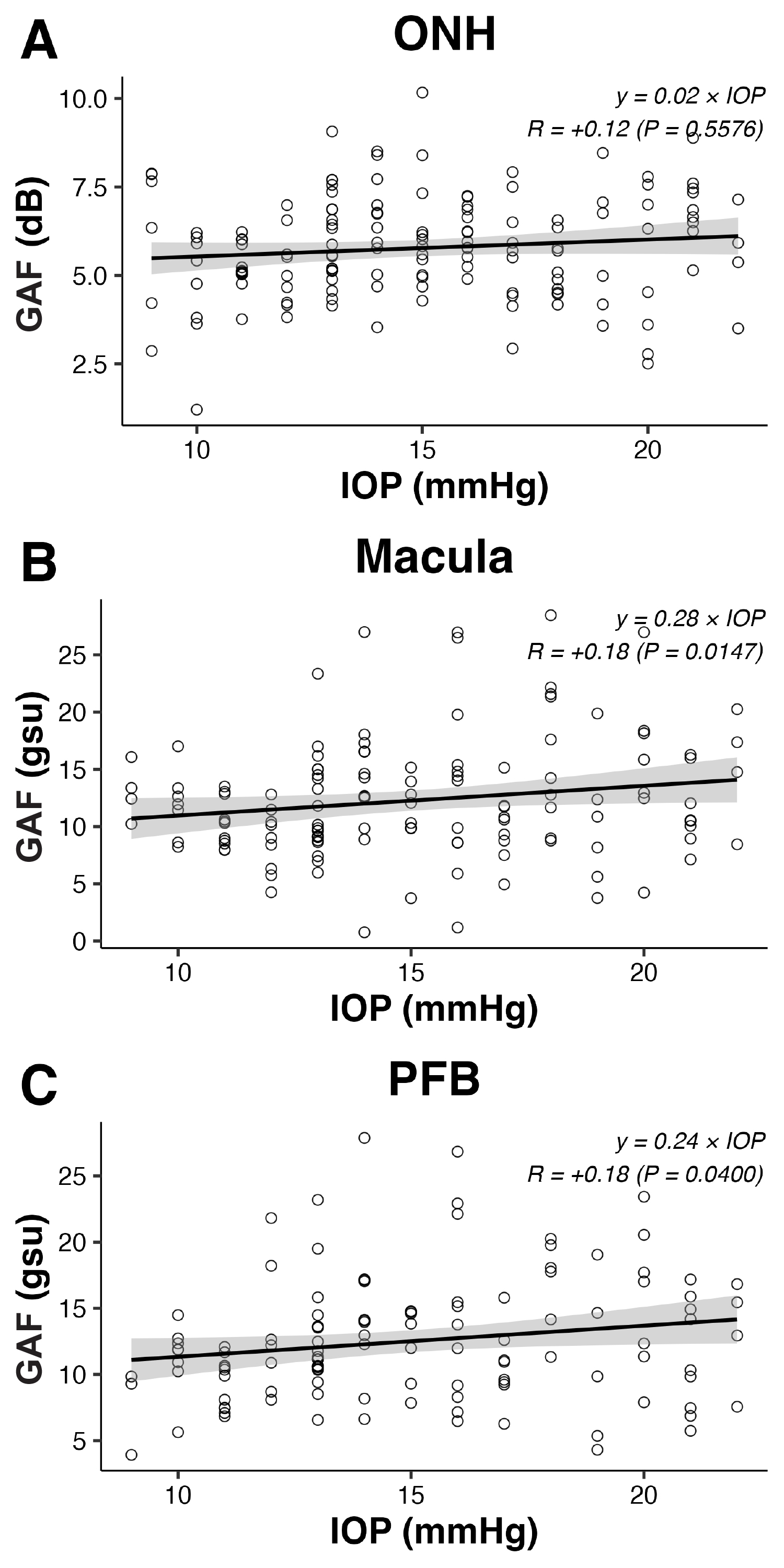

**Supplementary Figure S6. Association between intraocular pressure (IOP) and green autofluorescence (GAF) across the three retinal regions evaluated, excluding abnormal values. A**, ONH, GAF in decibels (dB); **B**, macula, GAF in grey scale units (gsu); **C**, PFB, GAF in grey scale units (gsu). Each data point represents one eye at a single imaging session with IOP recorded at the same visit. Solid lines show linear fits from linear mixed models (LMMs) accounting for intereye correlation and intraindividual repeated measures correlation, with 95 percent confidence bands. Panel annotations report the fitted equation together with the model R and *P* values.

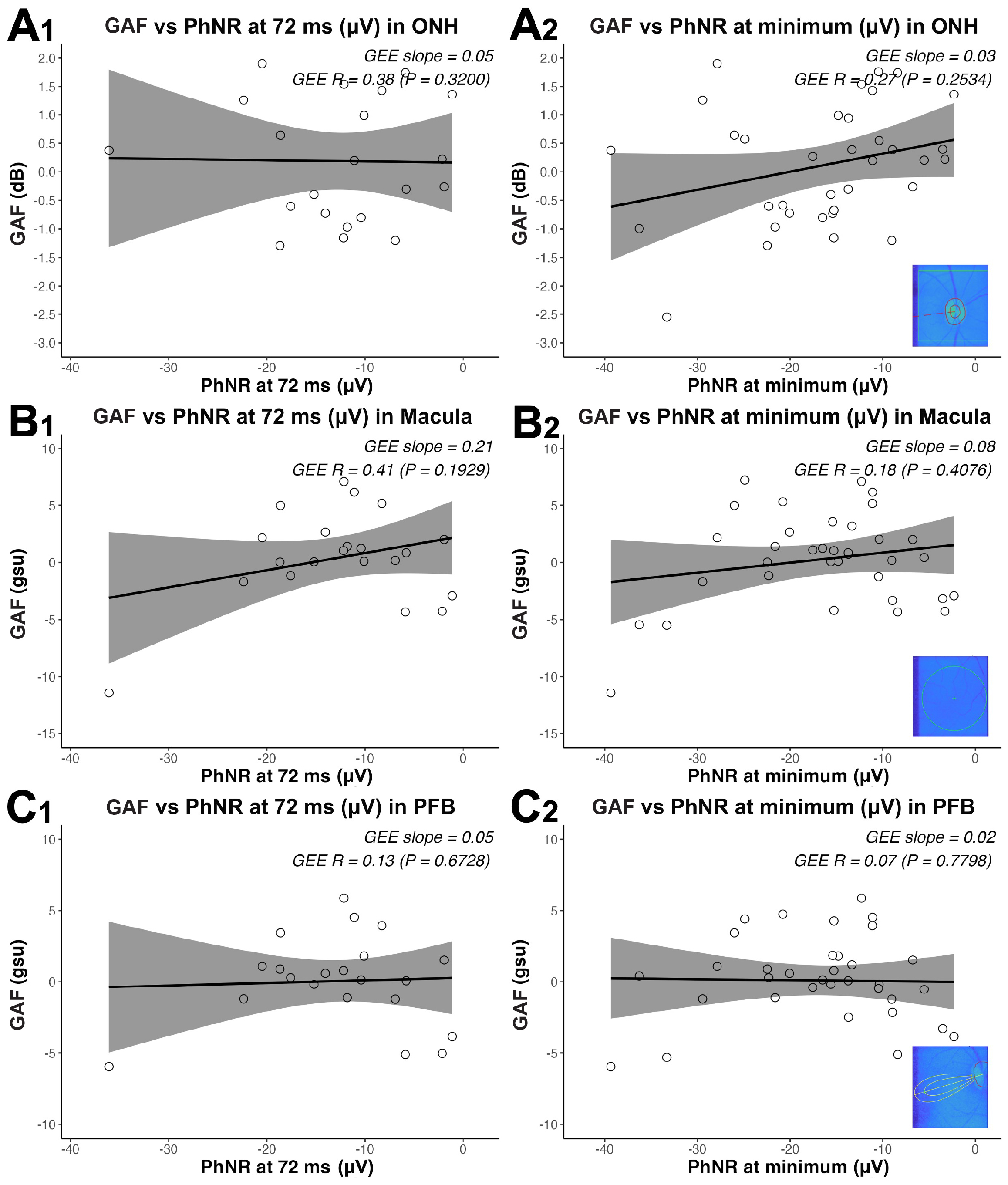

**Supplementary Figure S7. After adjusting for age, green autofluorescence (GAF) shows no significant association with photopic negative response PhNR amplitude at the optic nerve head (ONH), macula, or papillofoveal bundle (PFB). A1**, optic nerve head (ONH) GAF (dB) versus PhNR amplitude at 72 ms (µV); **A2**, ONH GAF (dB) versus PhNR amplitude at the minimum trough; **B1**, macula GAF (gsu) versus PhNR amplitude at 72 ms; **B2**, macula GAF (gsu) versus PhNR amplitude at the minimum trough; **C1**, papillofoveal bundle (PFB) GAF (gsu) versus PhNR amplitude at 72 ms; **C2**, PFB GAF (gsu) versus PhNR amplitude at the minimum trough. Each data point represents one eye at a single imaging session; PhNR testing and GAF imaging were performed within a maximum interval of three weeks. Solid lines show linear fits from mixed models accounting for intereye and intraindividual repeated measures correlation, with 95 percent confidence bands. Annotations in each panel report the slope (GAF units per µV), marginal R, and two-sided P. Insets show representative pseudocolor GAF images from the analyzed regions for reference.

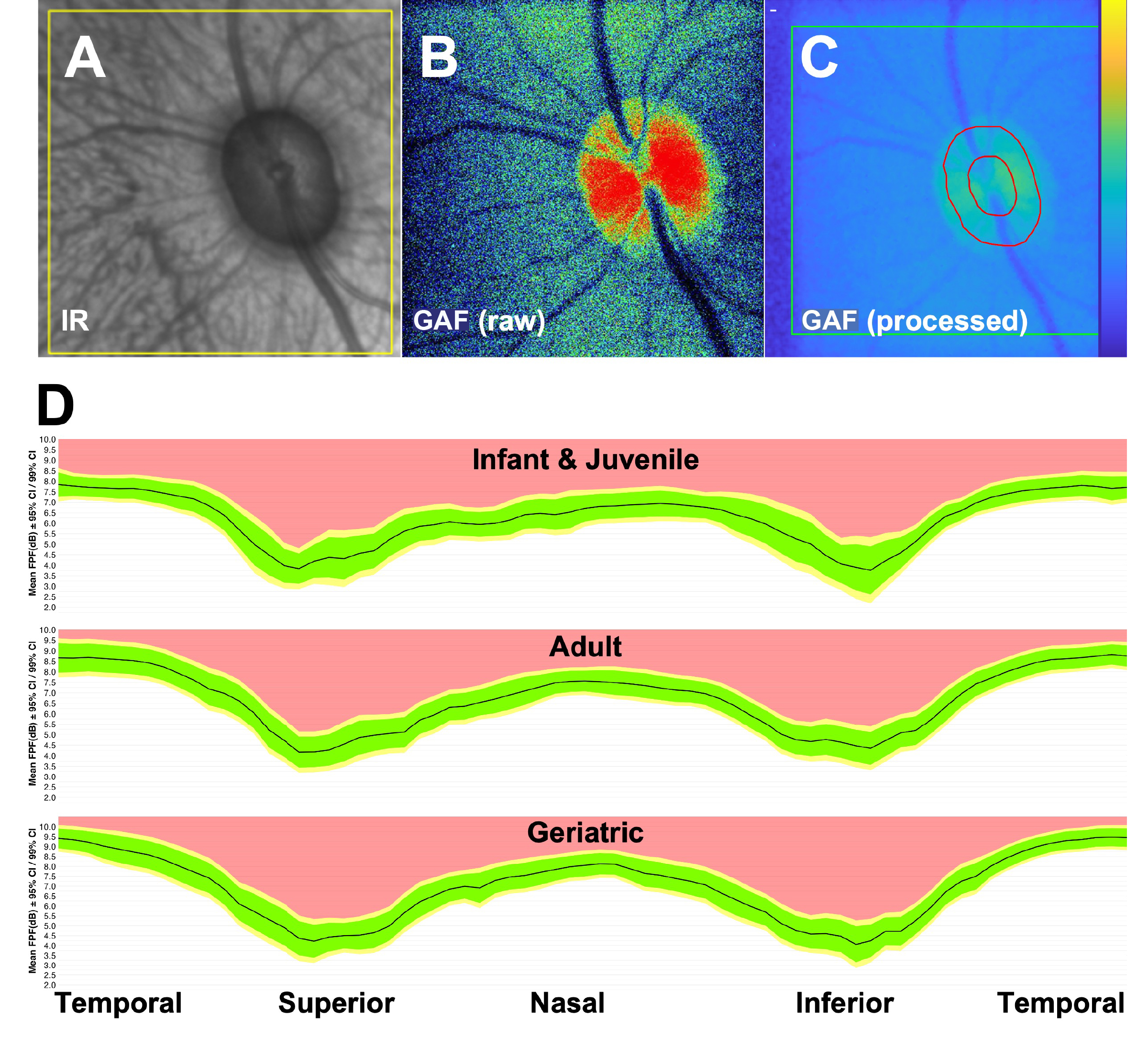

**Supplementary Figure S8. Standard workflow for optic nerve head green autofluorescence (GAF) imaging and age stratified circumferential profiles show a consistent pattern across ages: highest temporally, intermediate nasally, and lowest superiorly and inferiorly. A**, infrared reflectance image used to locate the field of view and confirm that the optic nerve head is centered and in focus. **B**, raw GAF frame from the same eye acquired immediately after centering on the optic nerve head. **C**, processed GAF map after background subtraction, flat field correction, and intensity normalization, with regions of interest delineated for quantitative analysis. **D**, mean peripapillary GAF profile (dB) around the optic nerve by age group defined as Infant (<1 year), Juvenile (1–5 years), Adult (>5–19 years), and Geriatric (>19 years); the black line represents the mean, the green dashed region denotes the 95 percent confidence interval, the yellow region denotes the 99 percent confidence interval, and values shown in red are considered abnormally increased.

### Supplementary Data

#### Supplementary Data S1. Axial length correction in the macular GAF scan.

Although axial length (AXL) values derived from published sources or external datasets cannot fully capture individual variability, we estimated the likely magnitude of the resulting scaling and magnification error across ages using both literature-based assumptions and a different ocular biometry dataset from the same colony but different animals (n = 209; age 0.2 to 31.6 years) that was used for independent studies [(Yiu et al. 2020)](https://sciwheel.com/work/citation?ids=10349361&pre=&suf=&sa=0&dbf=0). Published rhesus macaque ocular biometry supports a biphasic AXL trajectory. During early development, mean axial length is approximately 13.2 mm at birth and increases rapidly to about 18.6 to 19.2 mm by 4 to 5 years of age, approximately +0.33 mm per year [(Bradley et al. 1999; Qiao‑Grider et al. 2007; Yiu et al. 2020)](https://sciwheel.com/work/citation?ids=18475217,18475220,10349361&pre=&pre=&pre=&suf=&suf=&suf=&sa=0,0,0&dbf=0&dbf=0&dbf=0). After ocular maturation, AXL changes slowly, with an age slope of approximately +0.01 mm per year reported across a broad adult and geriatric age range [(Ross et al. 2025; Lin, Tran, Kim, Park, Chen, et al. 2021; Lin, Tran, Kim, Park, Stout, et al. 2021)](https://sciwheel.com/work/citation?ids=18475221,15710471,14847928&pre=&pre=&pre=&suf=&suf=&suf=&sa=0,0,0&dbf=0&dbf=0&dbf=0).

Therefore, the estimated scaling error across ages for the green autofluorescence (GAF) 21.5° × 17° macular field is as follows:

**Supplementary Table S9. Estimated retinal field size corresponding to the 21.5° × 17° GAF macular imaging field across age, based on colony axial length measurements.**

| **Age (y)** | **Literature based AXL (mm)** | **Estimated field (mm) V × H** | **Estimated area (mm²)** |
| --- | --- | --- | --- |
| 0.2 | 18.29 | 4.8 × 3.8 | 18.3 |
| 1 | 18.56 | 4.9 × 3.9 | 18.8 |
| 3 | 19.22 | 5.0 × 4.0 | 20.2 |
| 5 | 19.89 | 5.2 × 4.1 | 21.6 |
| 10 | 19.94 | 5.2 × 4.1 | 21.7 |
| 20 | 20.05 | 5.3 × 4.2 | 21.9 |
| 29 | 20.15 | 5.3 × 4.2 | 22.1 |

Notes: AXL, axial length; V, vertical; H, horizontal. Estimated field dimensions and areas represent the expected retinal sampling region corresponding to a fixed 21.5° × 17° imaging field at the listed axial lengths.

As expected, this corresponds to an estimated retinal field that increases as the eye grows, from approximately 4.8 × 3.8 mm (18.3 mm²) at 0.2 years to 5.3 × 4.2 mm (22.1 mm²) at 29 years for the fixed macular scan. We then applied an AXL based magnification correction to express macular GAF per standardized retinal area and refit the age association model (**Suppl. Fig. S9**). The raw macular GAF showed a strong positive association with age (slope = +0.52 gsu/year; **Suppl. Fig. S9A1**). After applying the AXL-based magnification correction, the age association remained significant but was modestly attenuated (slope = +0.453 gsu/year; *P* < 1×10⁻⁶; **Suppl. Fig. S9A2**), indicating that retinal scaling contributes only minimally to the overall age-related increase in macular signal. Future studies should aim at obtaining AXL at the same GAF imaging session to enable subject specific magnification correction.
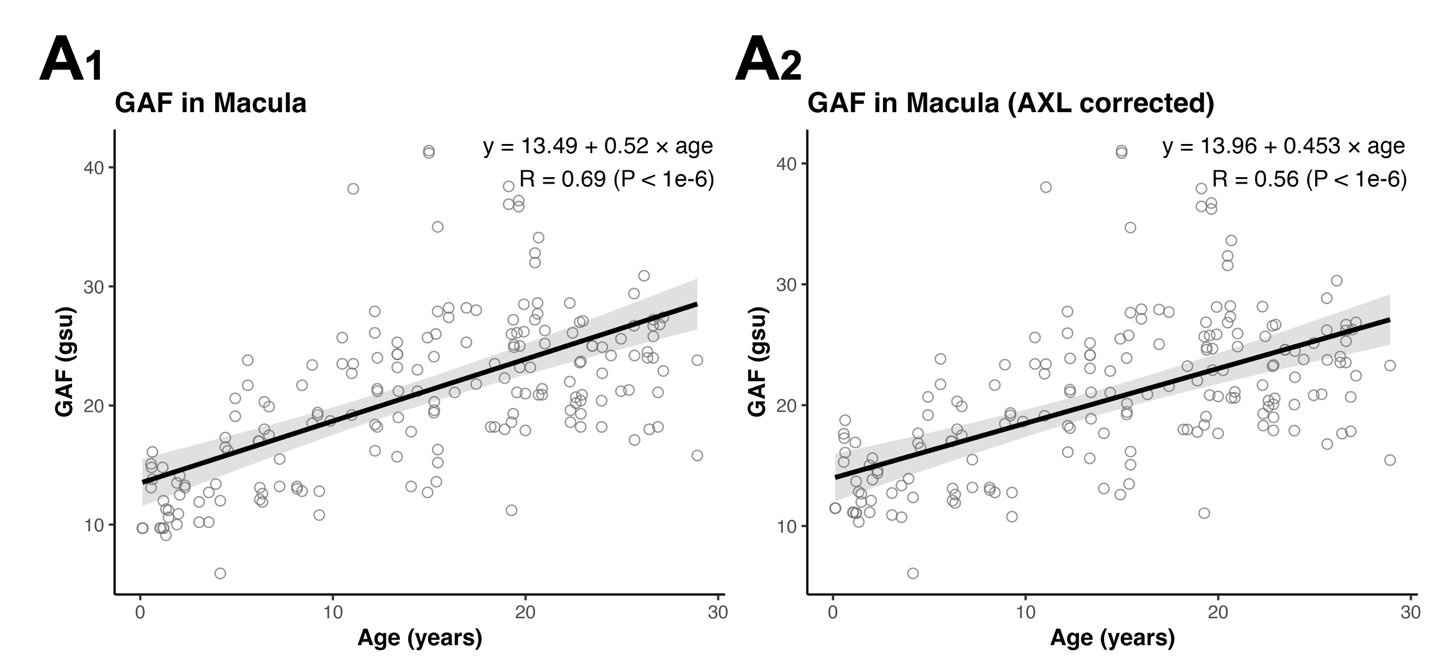

**Supplementary Figure S9. Analysis of the association between macular green autofluorescence (GAF) and age before and after correction for estimated ocular magnification.** **A1**, raw macular GAF (grey scale units, gsu) versus age (years) with linear regression fit (slope = +0.52 gsu/year, R = 0.69, *P* < 1×10⁻⁶). **A2**, macular GAF after normalization by the estimated retinal field area (based on age-matched colony axial length values) versus age with linear regression fit (slope = +0.453 gsu/year, R = 0.56, *P* < 1×10⁻⁶). Each point represents one eye at a single imaging session; shaded bands indicate 95% confidence intervals.
